# Unique metabolic strategies in Hadean analogues reveal hints for primordial physiology

**DOI:** 10.1101/2021.04.20.440570

**Authors:** Masaru Konishi Nobu, Ryosuke Nakai, Satoshi Tamazawa, Hiroshi Mori, Atsushi Toyoda, Akira Ijiri, Shino Suzuki, Ken Kurokawa, Yoichi Kamagata, Hideyuki Tamaki

## Abstract

Primordial microorganisms are postulated to have emerged in H_2_-rich alkaline Hadean serpentinite-hosted environments with homoacetogenesis as a core metabolism. However, investigation of two modern serpentinization-active analogues of early Earth reveals that conventional H_2_-/CO_2_-dependent homoacetogenesis is thermodynamically unfavorable *in situ* due to picomolar CO_2_ levels. Through metagenomics and thermodynamics, we discover unique taxa capable of metabolism adapted to the habitat. This included a novel deep-branching phylum, “*Ca*. Lithoacetigenota”, that exclusively inhabits Hadean analogues and harbors genes encoding alternative modes of H_2_-utilizing lithotrophy. Rather than CO_2_, these metabolisms utilize reduced carbon compounds detected *in situ* presumably serpentinization-derived: formate and glycine. The former employs a partial homoacetogenesis pathway and the latter a distinct pathway mediated by a rare selenoprotein – the glycine reductase. A survey of serpentinite-hosted system microbiomes shows that glycine reductases are diverse and nearly ubiquitous in Hadean analogues. “*Ca*. Lithoacetigenota” glycine reductases represent a basal lineage, suggesting that catabolic glycine reduction is an ancient bacterial innovation for gaining energy from geogenic H_2_ even under serpentinization-associated hyperalkaline, CO_2_-poor conditions. This draws remarkable parallels with ancestral archaeal H_2_-driven methyl-reducing methanogenesis recently proposed. Unique non-CO_2_-reducing metabolic strategies presented here may provide a new view into metabolisms that supported primordial life and the diversification of LUCA towards *Archaea* and *Bacteria*.

---

During the Hadean eon (~4.6-4.0 Ga), H_2_-rich hyperalkaline fluids generated from widespread serpentinization of ultramafic rocks are thought to have been conducive for the evolution of primordial life ^1–7^. Early microbial life is theorized to have catabolized H_2_ through homoacetogenesis ^4,8,9^, and recent studies point towards evolutionary antiquity of the central enzyme of the pathway, the bifunctional CO dehydrogenase/acetyl-CoA synthase or CODH/ACS ^10–12^. Our recent study also shows that an acetyl-CoA pathway-like chain of reactions can proceed in the presence of hydrothermal iron minerals ^13^, suggesting the pathway preceded life and life simply encapsulated this into cells ^14^. Protocells and the last universal common ancestor (LUCA) are hypothesized to have evolved within alkaline hydrothermal mineral deposits at the interface of serpentinization-derived fluid and ambient water (*e.g*., Hadean weakly acidic seawater) ^15–17^. Although such interfaces no longer exist (*i.e*., the Hadean Earth lacked O_2_ but most water bodies contain O_2_ on modern Earth), some anaerobic terrestrial and oceanic ecosystems harboring active serpentinization (*e.g*., Lost City hydrothermal field) ^18–25^ have been identified as modern analogues of ancient serpentinization-associated alkaline fluids. These are ideal ecosystems for investigating what kind of H_2_ catabolism may have supported early post-LUCA life venture away from the interface deeper into the hyperalkaline hydrothermal systems. Gaining independence from the gradient at the interface was likely a critical step in the evolution of life, and life likely headed towards the hyperalkaline fluids due to their reliance on serpentinization-derived energy sources and their cellular machinery being alkaliphilic (*i.e.*, moving towards the weakly acidic seawater would have been an unfavorable transition) ^5,15-17^. However, microbiologists have yet to provide insight into such metabolic strategies from extant organisms inhabiting modern analogues. In this study, we pair metagenomics and thermodynamics to characterize uncultured putative anaerobic H_2_ utilizers inhabiting alkaline H_2_-rich serpentinite-hosted systems (Hakuba Happo hot springs in Hakuba, Japan and The Cedars springs in California, USA; *p*H ~10.9 and ~11.9, respectively) ^26–28^ and elucidate lithotrophic catabolism that may have been relevant to early life.

## Thermogeochemistry

To evaluate whether homoacetogenesis is viable *in situ*, we examined the *in situ* geochemical environment and the thermodynamics of H_2_/formate utilization and homoacetogenesis. The spring waters of both Hakuba and The Cedars contained H_2_ (*e.g.*, 201-664 μM in Hakuba ^28^). Formate, another compound thought to be abiotically generated through serpentinization, was also detected in Hakuba (8 μM in drilling well #3 ^29^) and The Cedars (6.9 μM in GPS1). Acetate has also been detected *in situ* (4 μM in Hakuba ^29^ and 69.3 μM in The Cedars GPS1), suggesting these ecosystems may host novel H_2_- and/or formate-utilizing homoacetogens. Thermodynamic calculations confirm that H_2_ and formate are reductants *in situ* (*i.e*., H_2_ = 2H^+^ + 2e^-^ / Formate^-^ = H^+^ + CO_2_ + 2e^-^): the Gibbs free energy yields (ΔG) for oxidation (coupled with physiological electron carriers NADP^+^, NAD^+^, and ferredoxin) are less than −4.78 kJ per mol H_2_ and −24.92 kJ per mol formate in Hakuba, and −10.73 and −22.03 in The Cedars respectively (see Supplementary Results). However, serpentinite-hosted systems impose a unique challenge to homoacetogenesis – a key substrate, CO_2_, is at extremely low concentrations due to the high alkalinity. We estimate that the aqueous CO_2_ concentration is below 0.0004 nM in Hakuba (*p*H 10.7 and <0.1 μM TIC) and 0.003 nM in The Cedars (*p*H 11.9 and 35 μM TIC) ^26,28^. In Hakuba, H_2_/CO_2_-driven acetogenesis (ΔG of −1.68 kJ per mol acetate) cannot support microbial energy generation (ΔG ≤ −20 kJ per mol is necessary ^30^; Fig. S1). Moreover, in both Hakuba and The Cedars, one of the first steps in CO_2_-reducing homoacetogenesis, reduction of CO_2_ to formate, is unfavorable based on the thermodynamics presented above (ΔG > +24.92 or +22.02 kJ per mol formate). Thus, catabolic reduction of CO_2_ to acetate is thermodynamically challenging *in situ* and may only run if investing ATP (*e.g*., Calvin-Benson-Bassham cycle [−6 ATP; ΔG of −361.68 kJ per mol acetate in Hakuba] or reductive tricarboxylic acid [−1 ATP; −61.68 kJ per mol]). Under CO_2_ limitation, autotrophs are known to accelerate CO_2_ uptake through HCO_3_^-^ dehydration to CO_2_ (carbonic anhydrase) or carbonate mineral dissolution, but both only modify kinetics and are not effective in changing the maximum CO_2_ concentration (determined by equilibrium with carbonate species). In addition, in Hakuba, the CO_3_^2-^ concentration is too low (87.5 nM CO_3_^2-^) for carbonate mineral precipitation (*e.g.*, [CO_3_^2-^] must exceed 38.5 μM given K_s_ of 5×10^-9^ for CaCO_3_ and [Ca^2+^] of 0.13 mM).

Based on thermodynamic calculations, the energy obtainable from H_2_/CO_2_-driven homoacetogenesis is too small to support life in many Hadean analogues (Fig. S2a), yet acetate is detected in some of these ecosystems (Fig. S2b). Thus, CO_2_-independent electron-disposing metabolism may have been necessary for early life to gain energy from H_2_ in the hyperalkaline fluids of hydrothermal systems. Here, we explore the metabolic capacities of extant organisms living in CO_2_-limited Hadean analogues to gain insight into potential metabolic strategies that LUCA or early post-LUCA organisms beginning diversification towards *Bacteria* and *Archaea* may have utilized to thrive in alkaline serpentinite-active ecosystems.

## Diverse putative H_2_- and formate-utilizing organisms

Through metagenomic exploration of the two serpentinite-hosted systems (Table S1), we discover a plethora of phylogenetically novel organisms encoding genes for H_2_ and formate metabolism (19 bins with 73.2-94.8% completeness and 0.0-8.1% contamination [86.1% and 3.8% on average respectively]; available under NCBI BioProject PRJNA453100) despite challenges in acquisition of genomic DNA (15.7 and 18.9 ng of DNA from 233 and 720 L of filtered Hakuba Happo spring water, respectively; RNA was below the detection limit). We find metagenome-assembled genomes (MAGs) affiliated with lineages of *Firmicutes* (*e.g., Syntrophomonadaceae* and uncultured family SRB2), *Actinobacteria*, and candidate division NPL-UPA2 ^31^ (Fig. S3). We also recovered MAGs for a novel candidate phylum, herein referred to as “*Ca*. Lithoacetigenota”, that inhabits both Hakuba and The Cedars and, to our knowledge, no other ecosystems (Fig. 1a and S3). These genomes encode enzymes for oxidizing H_2_ and formate (*i.e*., hydrogenases and formate dehydrogenases ^32–39^; see Supplementary Results), suggesting that organisms *in situ* can employ H_2_ and formate as electron donors.

**Figure 1.**
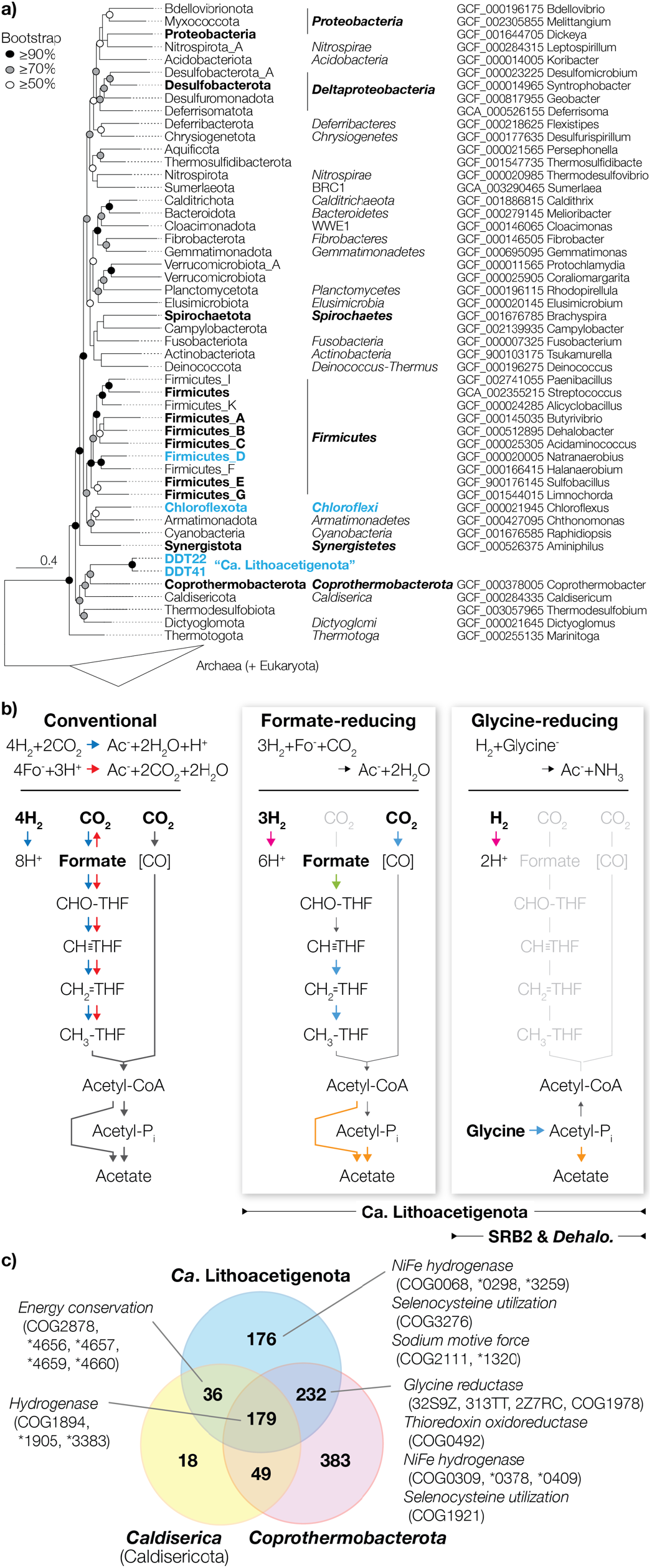
“*Ca*. Lithoacetigenota” phylogeny, lithotrophic acetate generation pathways, and comparative genomics with neighboring phyla. (a) A maximum likelihood tree was calculated using PhyML using the LG model, 4 gamma categories, and 100 bootstrap replicates from a concatenated alignment of universally conserved ribosomal protein sequences from representative genomes of individual phyla aligned with MAFFT (default parameters) and trimmed with trimAl prior to concatenation. Bootstrap values were recalculated using BOOSTER. Phyla that possess glycine reductases (black+bolded) and phyla for which glycine reductases were detected in serpentinite-hosted systems are indicated (blue+bolded). Phylum names are shown for both NCBI taxonomy (italicized) and GTDB classification. (b) Metabolic pathways potentially adapted to the CO_2_-limited hyperalkaline conditions encoded by “*Ca*. Lithoacetigenota” members and others: formate- and glycine-reducing acetate generation. Arrow colors indicate oxidative (pink), reductive (blue), ATP-yielding (orange), and ATP-consuming (green) steps. (c) Venn diagram of COGs/NOGs (as predicted by eggnog-mapper) fully conserved across all members of each phylum (genomes included in GTDB release 95 with completeness ≥85% and contamination ≤5%). COGs/NOGs related to lithotrophy and alkaliphily are highlighted. * “COG” abbreviated.

## “*Ca*. Lithoacetigenota” has unique site-adapted metabolism

Inspection of the serpentinite-hosted environment-exclusive phylum “*Ca*. Lithoacetigenota” reveals specialization to H_2_-driven lithotrophy potentially suitable for the low-CO_2_ *in situ* conditions (Fig. 1b). We discover that The Cedars-inhabiting population (*e.g.*, MAG BS5B28, 94.8% completeness and 2.9% contamination) harbors genes for H_2_ oxidation ([NiFe] hydrogenase Hox) and a nearly complete Wood-Ljungdahl pathway and an oxidoreductase often associated with acetogenesis – NADH:ferredoxin oxidoreductase Rnf ^40,41^ (Table S2 and S3). One critical enzyme, the formate dehydrogenase, is missing from all three “*Ca*. Lithoacetigenota” MAGs from The Cedars (and unbinned contigs), indicating that these bacteria can neither perform H_2_/CO_2_-driven nor formate-oxidizing acetogenesis (Fig. 1b). However, even without the formate dehydrogenase, the genes present can form a coherent pathway that uses formate rather than CO_2_ as a starting point for the “methyl branch” of the Wood-Ljungdahl pathway (*i.e.*, formate serves as an electron acceptor; Fig. 1b). This is a simple yet potentially effective strategy for performing homoacetogenesis while circumventing the unfavorable reduction of CO_2_ to formate. Coupling H_2_ oxidation with this formate-reducing pathway is thermodynamically viable as it halves the usage of CO_2_ (3H_2_ + Formate^-^ + CO_2_ = Acetate^-^ + 2H_2_O; ΔG of −29.62 kJ per mol acetate) and, as a pathway, is simply an intersection between the conventional H_2_/CO_2_-driven and formate-disproportionating acetogenesis (Fig. 1b). Although use of formate as an electron acceptor for formate-oxidizing acetogenesis is quite common, no previous homoacetogens have been observed to couple H_2_ oxidation with acetogenesis from formate, likely because CO_2_ has a much higher availability than formate in most ecosystems.

Interestingly, the Hakuba-inhabiting “*Ca*. Lithoacetigenota” (HKB210 and HKB111) also encodes Hox for H_2_ oxidation but lacks genes for homoacetogenesis (no homologs closely related to The Cedars population genes were detected even in unbinned metagenomic contigs). Perhaps this population forgoes the above H_2_/formate-driven homoacetogenesis because the estimated energy yield of the net reaction *in situ* (ΔG of −19.94 kJ per mol acetate) is extremely close to the thermodynamic threshold of microbial catabolism (slightly above −20 kJ per mol) and, depending on the actual threshold for “*Ca*. Lithoacetigenota” and/or even slight changes in the surrounding conditions (*e.g.*, ΔG increases by 1 kJ per mol if H_2_ decreases by 20 μM decreases in Hakuba), the metabolism may be unable to recover energy. Through searching the physicochemical environment for alternative exogenous electron acceptors and MAGs for electron-disposing pathways, we detected a low concentration of glycine *in situ* (5.4 ± 1.6 nM; Table S4) and found genes specific to catabolic glycine reduction (see next paragraph). We suspect that some portion of this glycine is likely geochemically generated *in situ*, given that (a) glycine is often detected as the most abundant amino acid produced by both natural and laboratory-based serpentinization (*e.g.*, H_2_ + Formate = Formaldehyde ⇒ Formaldehyde + NH_3_ = Glycine) ^1,7,42-47^ and (b) no other amino acid was consistently detectable (if glycine was cell-derived, other amino acids ought to also be consistently detected).

For utilization of the putatively abiotic glycine, the Hakuba “*Ca*. Lithoacetigenota” encode glycine reductases (Grd; Fig. 2 and S4; Tables S2 and S3) – a unidirectional selenoprotein for catabolic glycine reduction ^48,49^. Based on the genes available, this population likely specializes in coupling H_2_ oxidation and glycine reduction (Fig. 1b). Firstly, the genomes encode NADP-linked thioredoxin reductases (NADPH + Thioredoxin_ox_ à NADP^+^ + Thioredoxin_red_) that can bridge electron transfer from H_2_ oxidation (H_2_ + NADP à NADPH + H^+^) to glycine reduction (Glycine^-^ + Thioredoxin_red_ à Acetyl-P_i_ + NH_3_ + Thioredoxin_ox_). Secondly, though glycine reduction is typically coupled with amino acid oxidation (*i.e.*, Stickland reaction in *Firmicutes* and *Synergistetes* ^48,50^), similar metabolic couplings have been reported for some organisms (*i.e.*, formate-oxidizing glycine reduction [via Grd] ^51^ and H_2_-oxidizing trimethylglycine reduction [via Grd-related betaine reductase] ^52^). Thirdly, Grd is a rare catabolic enzyme, so far found in organisms that specialize in amino acid (or peptide) catabolism, many of which are reported to use glycine for the Stickland reaction (*e.g.*, *Peptoclostridium* of *Firmicutes* and *Aminobacterium* of *Synergistetes* ^53^). Lastly, the population lacks any discernable fermentative and respiratory electron disposal pathways and oxidative organotrophy (Table S2 and S3). Moreover, reflecting the lack of other catabolic pathways, the Hakuba “*Ca*. Lithoacetigenota” MAGs display extensive genome streamlining, comparable to that of *Aurantimicrobium* ^54,55^, “*Ca.* Pelagibacter” ^56^, and *Rhodoluna* ^57^ (Fig. S6). Thermodynamic calculations show that H_2_-oxidizing glycine reduction is thermodynamically favorable *in situ* (ΔG°’ of −70.37 kJ per mol glycine [ΔG of −85.84 in Hakuba]; Fig. S1). Further, based on the pathway identified, this metabolism is >10 times more efficient in recovering energy from H_2_ (1 mol ATP per mol H_2_) than acetogenesis utilizing H_2_/CO_2_ (0.075 mol ATP per mol H_2_ based on the pathway *Acetobacterium woodii* utilizes) or H_2_/formate (0.075 mol ATP per mol H_2_, assuming no energy recovery associated with the formate dehydrogenase). We also detect glycine reductases in The Cedars “*Ca*. Lithoacetigenota”, indicating that it may also perform this metabolism (ΔG of −76.87 in The Cedars, assuming 201 μM H_2_). Thus, we propose glycine as an overlooked thermodynamically and energetically favorable electron acceptor for H_2_ oxidation in serpentinite-hosted systems.

**Figure 2.**
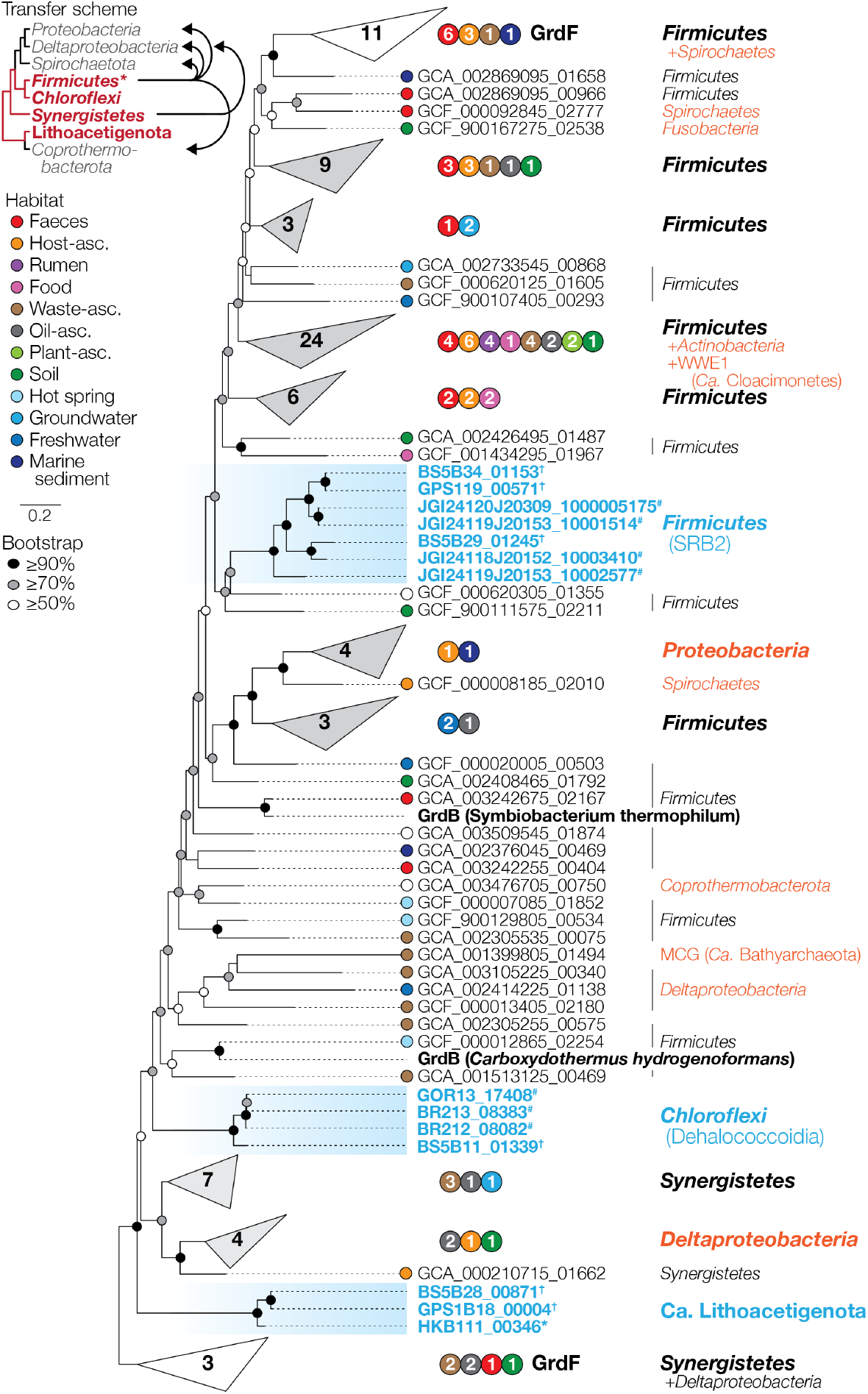
Phylogeny of serpentinite-hosted microbiome glycine reductase subunit GrdB homologs (Hakuba Happo hot spring*, The Cedars springs†, and other serpentinite-hosted system metagenomes#) and a brief scheme for evolutionary history of Grd. COG1978 homologs were collected from the representative species genomes in GTDB, filtered using a GrdB motif conserved across members of phyla known to perform glycine-reducing Stickland reaction and a GrdF motif conserved across sequences that form a distinct cluster around the biochemically characterized *Peptoclostridium acidaminophilum* sarcosine reductase subunit GrdF (see Methods and Supplementary Fig. S4), and clustered with 75% amino acid sequence similarity using CD-HIT (-c 0.75). A maximum likelihood tree was calculated as described in Fig. 1. For each sequence, the original habitat the isolate or MAG was obtained from is shown (white circle = unknown). Large sequence clusters were grouped (number of representative sequences included are shown). Note that the outgroup is a cluster of uncharacterized *Synergistetes* and *Deltaproteobacteria* sequences that was inferred to function as GrdF given that it shares the motif found in the *Firmicutes* GrdF. Taxa thought to have gained GrdB through horizontal transfer are shown in orange. See Supplementary Fig. S4 for complete tree. In the brief scheme of Grd evolution, the cladogram topology is based on Fig. 1a. Vertical transfer (red lines in cladogram) and horizontal transfer (black arrows) inferred from tree structures are shown. Phyla that may have acquired Grd vertically (red) and horizontally (gray) are indicated. GTDB phyla belonging to *Firmicutes* were grouped together.

Given the phylogenetic and metabolic uniqueness of these populations, we report provisional taxonomic assignment to “*Ca*. Lithoacetigenota” phylum. nov., “*Ca*. Thermoacetigena glycinireducens” gen. nov., sp. nov. (HKB111 and HKB210), and “*Ca*. Psychroacetigena formicireducens” gen. nov., sp. nov. (BS525, BS5B28, and GPS1B18) (see Supplementary Results). Based on a concatenated ribosomal protein tree, this serpentinite-hosted ecosystem-associated candidate phylum is closely related to a deep-branching group of bacterial phyla (Fig. 1a) (*e.g.*, *Caldiserica* [Caldisericota in GTDB phylogeny], *Coprothermobacterota*, and *Dictyoglomi* [Dictyoglomota] ^58^), many of which have extremely limited phylogenetic diversity (only 8, 1, and 2 genus-level lineages identified respectively via cultivation and metagenomics [based on GTDB release 95]) and ecological distribution on modern Earth. Comparative genomics shows that “*Ca*. Lithoacetigenota” shares 623 core functions (based on Bacteria-level COGs/NOGs predicted by eggnog-mapper shared by the two highest quality Hakuba and The Cedars MAGs HKB210 and BS5B28; Fig. 1c). When compared with the core functions of two closest related phyla (*Caldiserica* and *Coprothermobacterota*), 176 functions were unique to “*Ca*. Lithoacetigenota”, including those for NiFe hydrogenases (and their maturation proteins), selenocysteine utilization (essential for Grd), and sodium:proton antiporter for alkaliphily. With *Coprothermobacterota*, 232 functions were shared, including Grd, thioredoxin oxidoreductase (essential for electron transfer to Grd), and additional proteins for NiFe hydrogenases and selenocysteine utilization, pointing towards importance of H_2_ metabolism and glycine reduction for these closely related phyla. More importantly, among bacterial phyla in the deep-branching group, “*Ca*. Lithoacetigenota” represents the first lineage inhabiting hyperalkaliphilic serpentinite-hosted Hadean analogue ecosystems, suggesting that these organisms may be valuable extant windows into potential physiologies LUCA and/or early post-LUCA organisms may have taken (albeit with 4 billion years of evolution in between; see discussion regarding Grd below).

## Widespread glycine reduction in serpentinite-hosted systems

Uncultured members of *Chloroflexi* (Chloroflexota) class *Dehalococcoidia* inhabiting The Cedars and *Firmicutes* (Firmicutes_D) class SRB2 in Hakuba and The Cedars also possess glycine reductases (Table S3). In addition, these populations encode hydrogenases and formate dehydrogenases, suggesting that they may also link H_2_ and formate metabolism to glycine reduction. Moreover, we further discover closely related glycine reductases in other studied serpentinite-hosted systems (47~94% amino acid similarity in Tablelands, Voltri Massif, and Coast Range Ophiolite) ^18,19,24^. Phylogenetic analysis of the glycine-binding “protein B” subunit GrdB reveals close evolutionary relationships between glycine reductases from distant/remote sites (Fig. 2, S4, and S5). (Tablelands spring glycine reductase sequences were not included in the analysis as they were only detected in the unassembled metagenomic reads; 4460690.3; 69.7~82.2% similarity to Hakuba SRB2). Overall, “*Ca*. Lithoacetigenota”, *Dehalococcoidia*, and SRB2 glycine reductases are all detected in at least two out of the seven metagenomically investigated systems despite the diverse environmental conditions (*e.g.*, temperature), highlighting the potential importance of glycine metabolism by these clades in serpentinite-hosted systems. We suspect that glycine reduction may be a valuable catabolic strategy as the pathway requires few genes/proteins (a hydrogenase, Grd, acetate kinase, and thioredoxin oxidoreductase) and conveniently provides acetate, ammonia, and ATP as basic forms of carbon, nitrogen, and energy.

Comparison of the topology of the GrdB (Fig. 2, S4, and S5) and ribosomal protein (Fig. 1a) trees hints towards vertical transfer of GrdB dating back to one of the deepest divisions in the bacterial domain. Given the ecosystem specificity and deep phylogeny of the newly discovered phylum and Grd, catabolic glycine reduction may be a relatively ancient metabolism viable in serpentinization-related habitats, rapidly lost due to its low utility in modern ecosystems (*e.g.*, no severe nutrient/electron acceptor limitation and no excess glycine via abiotic generation), but repurposed by some anaerobes for the fermentative Stickland reaction in organic-rich ecosystems (*e.g.*, faeces and biodigesters) where excess amino acids are available but no favorable electron acceptors are accessible (*i.e.*, anaerobic). We identified the first archaeal GRD (in Miscellaneous Crenarchaeota Group [MCG] or *Ca.* Bathyarchaeota member BA-1; Fig. 2, S4, and S5), but phylogenetic analysis shows that the gene was gained through horizontal transfer and whether this gene truly belongs to this clade remains to be verified given that the source is a metagenome-assembled genome. Thus, the currently available data suggests that Grd (and catabolic glycine reduction) is an ancient bacterial innovation (*i.e.*, originated in *Bacteria*) developed post-LUCA and largely exclusive to the domain *Bacteria*. Further investigation of Hadean analogues is necessary to verify the history of Grd (still unclear based on Bayesian inference of GrdB phylogeny; Fig. S5), uncover other basal Grd and explore whether glycine reduction (or Grd family proteins) can be dated back further (more ancient *Bacteria* or LUCA; *i.e.*, necessary to find more basal bacterial and/or archaeal homologs).

## Other characteristics of putative indigenous homoacetogens

In contrast with members of “*Ca*. Lithoacetigenota”, several other putative homoacetogenic populations encode the complete Wood-Ljundgahl pathway (Table S2 and S3), indicating that other forms of acetogenesis may also be viable *in situ*. One putative homoacetogen in The Cedars, NPL-UPA2, lacks hydrogenases but encodes formate dehydrogenases. Although the NPL-UPA2 population cannot perform H_2_/formate-driven acetogenesis, it may couple formate oxidation with formate-reducing acetogenesis – another thermodynamically viable metabolism (ΔG of −50.90 kJ per mol acetate in The Cedars; Fig. 1b). The pathway uses CO_2_ as a substrate but has lower CO_2_ consumption compared to H_2_/CO_2_ homoacetogenesis and can produce intracellular CO_2_ from formate. In Hakuba, an *Actinobacteria* population affiliated with the uncultured class UBA1414 (MAG HKB206) encodes hydrogenases and a complete Wood-Ljungdahl pathway (Table S3) and, thus, may be capable of H_2_/formate or the above formate-disproportionating acetogenesis (Fig. 1b). Indeed, the UBA1414 population was enriched in Hakuba-derived cultures aiming to enrich acetogens using the H_2_ generated by the metallic iron-water reaction ^59^ (Fig. S7). Many populations encoding a complete Wood-Ljungdahl pathway possess monomeric CO dehydrogenases (CooS unassociated with CODH/ACS subunits; NPL-UPA2, *Actinobacteria, Syntrophomonadaceae* [Hakuba and The Cedars], and *Dehalococcoidia* [The Cedars]; Table S2). Although CO is below the detection limit in Hakuba (personal communication with permission from Dr. Konomi Suda), another study shows that CO metabolism takes place in an actively serpentinizing system with no detectable CO ^60^. Given that CO is a known product of serpentinization ^24,60^, it may be an important substrate for thermodynamically favorable acetogenesis *in situ*. However, further investigation is necessary to verify this.

Another interesting adaptation observed for all putative homoacetogens detected in Hakuba and The Cedars was possession of an unusual CODH/ACS complex. Although *Bacteria* and *Archaea* are known to encode structurally distinct forms of CODH/ACS (designated as Acs and Cdh respectively for this study), all studied Hakuba/The Cedars putative homoacetogens encode genes for a hybrid CODH/ACS that integrate archaeal subunits for the CO dehydrogenase (AcsA replaced with CdhAB) and acetyl-CoA synthase (AcsB replaced with CdhC) and bacterial subunits for the corrinoid protein and methyltransferase components (AcsCDE) (Table S2). The *Firmicutes* lineages also additionally encode the conventional bacterial AcsABCDE. Given that all of the identified putative homoacetogens encode this peculiar hybrid complex, we suspect that such CODH/ACS’s may have features that adapted to the high-*p*H low-CO_2_ conditions (*e.g.*, high affinity for CO_2_ and/or CO). In agreement, a similar hybrid CODH/ACS has also been found in “*Ca*. Desulforudis audaxviator” inhabiting an alkaline (*p*H 9.3) deep subsurface environment with a low CO_2_ concentration (below detection limit ^61,62^).

## Implications for primordial biology

Based on one of the most plausible theories of the origin of life, LUCA inhabited alkaline geochemically active sites ^63^, like serpentinite-hosted systems widespread across Earth during the Hadean to early Archean. Although reconstruction of the LUCA’s physiology is challenging ^64^, energy acquisition through H_2_-oxidation-driven CO_2_ reduction to acetyl-CoA (*i.e.*, H_2_/CO_2_ homoacetogenesis) is theorized to be a core feature of primordial metabolism ^11^. However, the free energy yield of such homoacetogenesis decreases with CO_2_ limitation and increasing temperature (Fig. S2a), suggesting that LUCA and/or early post-LUCA organisms may have encountered CO_2_-related thermodynamic challenges. Another study also points out that CO_2_ speciation towards a less bioavailable form (carbonate) under hyperalkaline pH may have been a potential problem for primordial life ^65^. At the interface of the alkaline fluids and seawater where life is thought to have originated ^7,15-17^, H_2_ and CO_2_ from the respective fluids could have come in contact and allow CO_2_-dependent H_2_ utilization; however, CO_2_-dependent metabolism would have become thermodynamically, and perhaps kinetically ^66^, challenging as early post-LUCA organisms ventured away from the interface and deeper into serpentinite-hosted systems. Through the investigation of CO_2_-poor Hadean analogues, we discover a novel Hadean analogue-exclusive alkaliphilic phylum that belongs to a deep-branching group of bacterial phyla and possesses unconventional thermodynamically favorable less CO_2_-dependent H_2_-oxidizing metabolic strategies (*e.g.*, coupled with formate or glycine reduction) that may be compelling candidates for early H_2_-dependent lithotrophy.

The thermodynamic and energetic favorability of catabolic H_2_-oxidizing glycine reduction mediated by GRD makes it a competitive metabolism for primordial *Bacteria*. Catabolic H_2_-utilizing glycine reduction also draws remarkable parallels with homoacetogenesis and ancestral archaeal metabolism. Like homoacetogenesis, glycine reduction has a rare feature thought to be essential for primordial metabolism – the ability to reductively synthesize acyl-phosphate for substrate-level phosphorylation in the cytosol ^67–70^ – indicating high utility for ancient organisms. Glycine reduction also shares features with a metabolism recently proposed to be ancestral in *Archaea* – H_2_-oxidizing methyl-reducing methanogenesis ^71,72^. Both are (nearly) domain-exclusive, potentially primordial, CO_2_-independent, and energetically more efficient than H_2_/CO_2_-driven acetogenesis (in terms of energy recovered per mol H_2_). Moreover, both utilize reduced serpentinization-derived carbon compounds, a thiol/disulfide electron carrier as an electron donor, and NiFe hydrogenases as an upstream electron source (Fig. 3). It is tempting to speculate that early post-LUCA *Bacteria* and *Archaea* both integrated geogenic thermodynamically favorable electron acceptors to adapt to thrive under the extreme conditions and interestingly occupied non-competing niches by using different acceptors (glycine and methyl compounds respectively) and energy acquisition strategies (substrate-level phosphorylation and proton motive force respectively), whereby allowing the two to co-exist with little competition. If true, we may begin to speculate that, as *Bacteria* and *Archaea* weaned themselves from serpentinite-hosted systems and explored the non-alkaline habitats of Earth, GRD and MCR were connected to the methyl branch of the Wood-Ljungdahl pathway and repurposed to facilitate CO_2_ fixation ^73^ and CO_2_-reducing methanogenesis respectively, both CO_2_-dependent pathways relevant today. However, further investigations are necessary to verify because, unlike MCR, information on GRD is limited: glycine-mediated autotrophy was only discovered recently, organisms still using GRD for catabolic glycine reduction are low in diversity on modern Earth given the abundant availability of other electron acceptors, and the geobiology of glycine-generating serpentinite-hosted systems have not been studied extensively.

**Figure 3.**
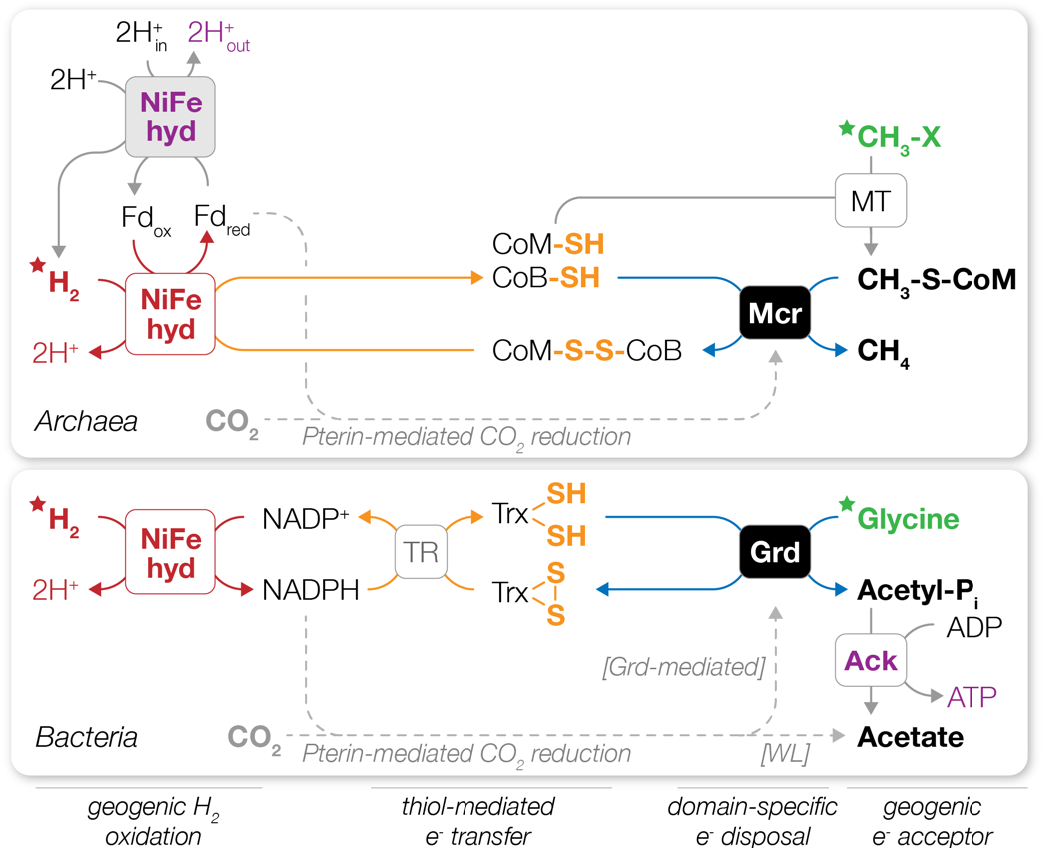
Proposed non-CO_2_-reducing metabolic strategies of ancestral archaea and bacteria inhabiting serpentinite-hosted systems. Geogenic carbon compounds, in particular methyl groups for archaea and glycine for bacteria (marked by a green star), can serve as electron acceptors for the oxidation of geogenic H_2_ (red star). The bacterial glycine-reducing H_2_-dependent lithotrophy is strategically similar to the ancestral archaeal methyl-reducing methanogenesis recently suggested ^71,72^. Both reactions can be mediated by nickel-iron [NiFe] hydrogenase H_2_ oxidation (red) and reduction of thiol/disulfide electron carriers (orange) and forgo pterin-mediated CO_2_ reduction (gray dotted line). Energy recovery (purple) is contrasting between the two metabolisms: archaea employing proton motive force (membrane-bound proteins gray) and bacteria substrate level phosphorylation. Reactions are not balanced with H^+^, H_2_O, and P_i_. The pathways shown are based on “*Ca*. Methanofastidiosum” ^74^ and “*Ca*. Thermolithoacetigena glycinireducens”. NiFe hyd, NiFe hydrogenase; Fd, ferredoxin; CoM-SH, coenzyme M thiol; CoB-SH, coenzyme B thiol; CoM-S-S-CoB, heterodisulfide of coenzyme M and coenzyme B; Mcr, methyl-coenzyme M reductase; CH_3_-X, methyl compounds; MT, methyltransferase; CH_3_-S-CoM, methyl-coenzyme M; TR, thioredoxin reductase; Trx-(SH)_2_, thioredoxin dithiol; Trx-S_2_, thioredoxin disulfide; Grd, glycine reductase; acetyl-P_i_, acetyl-phosphate; Ack, acetate kinase.

As was the case for MCR, metagenomic exploration of the uncharted rare biosphere provided key data for genes and core pathways that provide a glimpse into how primordial life or early bacteria may have proliferated (though we must still take caution in interpretation given the uncertainty associated with metagenomics). The thermodynamic (i.e., more favorable than the CO_2_-reducing counterpart) and evolutionary (i.e., ancestral) parallels between archaeal methyl-reducing and bacterial glycine-reducing H_2_-dependent lithotrophy warrant investigation of whether these metabolisms did indeed play a role in the formation or fixation of the two domains. Further studies on the evolutionary origin, antiquity, and history of glycine reduction will shed light on primordial way of life as well as bacterial/prokaryotic diversification.

## Supporting information

Supplementary Results, Figures S1-S7, and Tables S1-S4

## Author contribution

MKN and RN designed and performed metagenomic, phylogenetic, and thermodynamic analyses, interpreted the data, and wrote the manuscript. ST performed sampling, DNA extraction, and cultivation. HM, AT, and KK performed metagenomic sequencing and metagenomic analysis. AI performed chemical measurements. SS supported chemical data collection. HT and YK designed the project, supervised sampling and cultivation, and supported manuscript preparation.

## Acknowledgements

The authors greatly appreciate Mr. Sejima and others of Happo-one Development Co., Ltd. for their kind permission and cooperation in conducting field studies at Hakuba Happo hot springs. This work was supported by the JSPS KAKENHI Grant-in-Aid for Scientific Research on Innovative Areas, “Hadean Bioscience” project (no. JP26106006). This work was partly funded by the JSPS KAKENHI Grant-in-aid for Scientific Research nos. JP26710012 and JP18H02426 to H. Tamaki, JP15H05620 to R. Nakai, and JP18H03367 to M.K. Nobu.

## Competing interests

The authors declare no competing interests.

## Methods

### Sampling site and sample collection

The Hakuba Happo samples for geochemical and microbiological analysis were artificially pumped from a drilling well (700 m in depth), which was previously described and named Happo #3 (36° 42’ N 137° 48’ E ^28^). For microbiological analysis, two spring water samples were taken at different time points, 233 L taken in July 2016 (labelled HKB701) and 720 L taken in October 2016 (labelled HKB702), respectively. To collect cells, samples were filtered through a 0.1-μm Omnipore™ membrane filter (Merck Millipore) using a 90 mm diameter stainless-steel filter holder (Merck Millipore) attached to FDA Viton^®^ tubing (Masterflex) at a sampling site. After filtration, filters were immediately transferred to sterile tubes and frozen in a dry ice-ethanol bath, transported in dry ice, and stored at −80°C until DNA extraction. For NH_3_ and amino acid analysis, water samples were collected in October 2017, filtered as described above, transferred to dry-heat-sterilized nitrogen-purged 100-ml glass vials, and stored at 4°C.

### Geochemical analysis

The water temperature of hot spring water was measured using a thermometer (CT-430WP, Custom Ltd.) at a site. The pH, oxidation reduction potential (ORP), electrical conductivity (EC), and dissolved oxygen (DO) level were determined with portable devices, including a *p*H meter (D-23, Horiba), an ORP meter (RM-30P, TOA-DKK), an EC meter (CM-31P, TOA-DKK), and a DO meter (DO-31P, TOA-DKK), correspondingly. The ion concentrations of Na^+^, K^+^, Ca^2+^, and NO_3_^-^ were determined using portable sensors (LAQUAtwin™ series, Horiba). The *in situ* NH_3_ concentration was determined by measuring aqueous NH_4_^+^ and gaseous NH_3_ (purged with N_2_ gas, gas dissolved into deionized water, and measured dissolved NH_4_^+^) of a sample stored as described above using high-performance liquid chromatography (HPLC; Prominence; Shimadzu), then adding the two together. For amino acid quantification, the sample was concentrated under a stream of nitrogen gas and then analyzed following Shimadzu protocol no. L323 (https://www.ssi.shimadzu.com/products/literature/lc/L323.pdf) using HPLC with minor modifications (fluorescence detector RF-20Axs; sodium hypochlorite solution was not added for detection of proline). The Cedars spring concentrations of formate and acetate were determined by Isotope-Ratio-Monitoring Liquid Chromatography Mass Spectrometry (IRM-LCMS); Thermo-Finnigan Delta Plus XP isotope-ratio mass spectrometer connected to LC IsoLink, as described by Heuer et al. ^75^ and Ijiri et al. ^76^.

### Thermodynamic calculations

The Hakuba and Cedars calculations Gibbs free energy yield (ΔG) are based on ΔG°_f_ and ΔH°*f* values at 298 K, respective *p*H (10.8 and 11.9), and adjustment to the *in situ* temperatures (48 and 17°C) through the Gibbs-Helmholtz equation ^77^. The effect of pressure was approximated as described by Wang *et al* ^78^. For both Hakuba and The Cedars calculations, the glycine concentration (5.4 nM) was based on measurements from Hakuba. Formate, acetate, and NH_3_ concentrations were based on respective measurements from Hakuba (8 μM formate, 4 μM acetate, and 2.9 μM NH_3_) and The Cedars (6.9 μM formate [average of 6.777 and 7.079 μM measured on September 2017], 69.3 μM acetate [average of 69.601 and 68.967 μM measured on September 2017], and 1 μM NH_3_ [below detection limit]). For Hakuba, the H_2_ concentration measured in Hakuba drilling well #3 (DNA source) was used (201 μM H_2_). For The Cedars, the highest detected H_2_ concentration in Hakuba was used (664 μM H_2_ in drilling well #1).

### Metagenome sequencing, assembly and binning

The filter was aseptically cut into 16 equal pieces using sterilized tweezers, and each piece was placed in the bead-beating tube (Lysing Matrix E tube; MP Biomedicals). After DNA extraction following the bead-beating method described previously ^79^, the 16 DNA samples were mixed and then stored at −80°C until used. Sequence libraries were prepared with Nextera XT DNA Library Preparation kit (Illumina) with a genomic DNA fragment size ranging from 200 to 2,000 bp. These libraries sequenced on HiSeq2500 sequencing platform (Illumina) with HiSeq Rapid SBS kit v2 (Illumina), generating paired-end reads up to 250 bp. The generated sequences were trimmed using Trimmomatic v0.33 ^80^ with a quality cutoff of 30, sliding window of 6 bp, and minimum length cutoff of 78 bp. The trimmed sequences were assembled using SPAdes v3.10.1 ^81^ with the “-meta” option and k-mer values of 21, 31, 41, 53, 65, and 77. The assembled contigs were binned using MaxBin2.2.1 ^82,83^. The completeness and contamination of each bin was checked using CheckM ^84^. These bins were manually curated as described in our metagenomics study ^40^. Genes were then annotated using Prokka v1.12 ^85^ and eggnog-mapper ^86^. For interpretation and comparison of microbial metabolism, bin genomes were also constructed from public metagenomic data generated from The Cedars ^20^ (trimmed with sliding window of 6, quality cutoff of 20, and minimum length of 68 bp through Trimmomatic v0.33, normalized using BBMap 36.99 (https://jgi.doe.gov/data-and-tools/bbtools/) with target and minimum coverages of 40 and 2, assembled using SPAdes v3.10.1 with the “-meta” option and k-mer values of 21, 33, 45, 55, 67, and binned through MaxBin2.2.1) and were then analyzed collectively.

### Phylogenomic and phylogenetic analysis

For tree construction, sequences were aligned with MAFFT ^87^ v7.453 (default parameters) and trimmed using trimAl ^88^ v1.2rev59 (-gt 0.9). For ribosomal protein trees, a concataenated alignment of universally conserved ribosomal proteins ^89^ was used. Protein sequences were retrieved by downloading the GTDB ^90^ database and predicting protein sequences using Prokka ^85^ 1.14 (--kingdom Bacteria/Archaea --rnammer --addgenes --mincontiglen 200). Maximum likelihood trees were calculated using PhyML ^91^ v3.3.20190909 using the LG ^92^ model and 100 bootstrap replicates (-b 100 -d aa -m LG -v e). Bootstrap values were recalculated using BOOSTER ^93^. Sequence clustering was performing through CD-HIT ^94^ v4.8.1. For glycine reductase GrdB and sarcosine reductase GrdF, conserved motifs were predicted by first identifying fully conserved residues in the sequence cluster including the biochemically characterized *Peptoclostridium acidaminophilum* GrdF (see Supplementary Fig. S4; YxNx(6)GGE x(34,38) CGD x(27,35) GPxF[NF]AGRYG x(150,181) IHGGYDRx(6)[IP]x(4)PxD x(19,20) TTGTGTx(7)F x(12) [HILV]), then identifying fully conserved residues in the phylogenetic clusters (see Supplementary Fig. S4) that include GrdB from phyla known to perform the Stickland reaction (*Firmicutes, Spirochaetes*, and *Synergistetes*) subtracting any sequences clusters that contain the GrdF motif above (YxNx(6)GGE x(34,38) CGD x(27,35) GPxF[NF]AGRYG x(157,178) AHGGxD[QTAP] x(8) RV[IL]PxD x(19,20) TxGNxTxV). Bayesian trees were calculated using Phylobayes (MPI) v1.8 using the LG model with 4 gamma categories (-lg -ncat 1 -dgam 4) with four chains (chain length 10000; -x 1 10000) of which three chains that converged (maxdiff=0.1859, meandiff=0.014) were used for determination of a consensus tree with a burn-in of 1000 (nodes with posterior probability less than 0.8 were collapsed; bpcomp -c 0.8 -x 1000 1).

### Data availability

The datasets generated during and/or analyzed during the current study are available in the National Center for Biotechnology Information (NCBI) under BioProject PRJNA453100 and BioSamples SAMN08978938-SAMN08978962.

## Notes

### Competing Interest Statement

The authors have declared no competing interest.

