## Supplementary Results, Figures S1-S7, and Tables S1-S4 for "Unique metabolic strategies in Hadean analogues reveal hints for primordial physiology"

- Supplementary Information -

Masaru Konishi Nobu[1†*], Ryosuke Nakai[1,2†], Satoshi Tamazawa[1,3], Hiroshi Mori[4], Atsushi Toyoda[4], Akira Ijiri[5], Shino Suzuki[6,7], Ken Kurokawa[4], Yoichi Kamagata[1], and Hideyuki Tamaki[1*]

Affiliation:
[1] Bioproduction Research Institute, National Institute of Advanced Industrial Science and Technology (AIST), 1-1-1 Higashi, Tsukuba, Ibaraki 305-8566, Japan
[2] Bioproduction Research Institute, National Institute of Advanced Industrial Science and Technology (AIST), 2-17-2-1, Tsukisamu-Higashi, Sapporo, 062-8517, Japan
[3] Horonobe Research Institute for the Subsurface Environment (H-RISE), Northern Advancement Center for Science & Technology, 5-3 Sakaemachi, Horonobe, Teshio, Hokkaido, 098-3221, Japan
[4] National Institute of Genetics, 1111 Yata, Mishima, Shizuoka 411-8540, Japan
[5] Kochi Institute for Core Sample Research, Japan Agency for Marine-Earth Science and Technology (JAMSTEC), 200 Monobe Otsu, Nankoku, Kochi, Japan
[6] Institute for Extra-cutting-edge Science and Technology Avant-garde Research (X-star), JAMSTEC, Natsushima 2-15, Yokosuka, Kanagawa 237-0061, Japan
[7] Institute of Space and Astronautical Science (ISAS), Japan Aerospace Exploration Agency (JAXA), 3-1-1 Yoshinodai, Chuo-ku, Sagamihara, Kanagawa 252-5210, Japan

[†] These authors contributed equally.

### Table of Contents

Supplementary Results ..................... 2

Supplementary Figures ..................... 3
- Figure S1 3
- Figure S2 4
- Figure S3 5
- Figure S4 6
- Figure S5 7
- Figure S6 8
- Figure S7 9
- References 10

Supplementary Tables ..................... 11
- Table S1 11
- Table S2 12
- Table S3 13
- Table S4 14

### $H_2$ and formate metabolism

Assuming that the hydrogenases and formate dehydrogenases *in situ* use NADP(H) or NAD(H)+ferredoxin (*i.e.*, electron-bifurcating) (an assumption confirmed based on analysis of the metagenome-assembled genomes we recover; see below), $H_2$ and formate are likely reductants. In Hakuba, we estimate ΔG of +8.64 and +4.78 kJ per mol $H_2$ for $H_2$ generation through the respective pathways, assuming (i) literature cytosolic electron carrier redox potentials (-370 mV for NADPH, -320 mV NADH, and -450 mV Fd), (ii) cytosolic pH of 8.8 (two units lower than extracellular milieu [1]), and (iii) intracellular $H_2$ concentrations similar to surrounding environment. As for formate metabolism, formate dehydrogenases are predicted to run in the oxidative direction because $CO_2$ reduction to formate is also endergonic in situ (ΔG of +30.28 and +24.92 kJ per mol formate depending on the electron carrier, with identical assumptions). $H_2$ and formate generation can only become exergonic ($\Delta G > 0$) if cytosolic $H_2$ and formate reach below 266 nM and 0.115 nM respectively. Similarly, in The Cedars (estimated cytosolic pH of 9.9), $H_2$ and formate must be less than 20.6 nM and 5.18 nM respectively.

For $H_2$ metabolism, we identify putative NADP-reducing hydrogenases (HoxEFUHY in *Actinobacteria* and "*Ca*. Lithoacetigenota" and HndABCD in Firmicutes) and NAD/Fd-dependent electron-confurcating hydrogenases (HydABC in Firmicutes) [2-9]. HoxEFUHY typically uses NAD(H) as an electron carrier, but the Hox-related hydrogenases of HKB210 and BS525 consistently associate with a sixth subunit containing a putative NADPH-binding GltD domain, suggesting that these hydrogenases may employ NADP(H) as an electron carrier rather than NAD(H), a phenomenon that has also been reported for the *Ralstonia eutropha* HoxEFUHYI. For formate metabolism, we predict NADP-dependent formate dehydrogenases in one *Syntrophomonadaceae* population and putative electron-confurcating formate dehydrogenases in *Actinobacteria*, NPL-UPA2, and a *Syntrophomonadaceae* population.

### Nomenclature

Lithoacetigenota
Lithoacetigena (Li.tho.a.ce'ti.ge'na. Gr. n. *lithos* stone; N.L. n. *acidum aceticum* acetic acid; L. suff. - *genus -a - um* (from L. v. *gigno*) producing acetate; N.L. n. *Lithoacetigena* producing acetate from inorganic substrate).
Thermoacetigena (Ther.mo.a.ce'ti.ge'na. Gr. adj. *thermos* hot; N.L. n. *acidum aceticum* acetic acid; L. suff. -*genus -a -um* (from L. v. *gigno*) producing acetate; N.L. n. *Thermoacetigena* producing acetate under thermophilic conditions). Thermoacetigena glycinireducens (gly.ci.ni.re.du'cens. N.L. glycinum glycine; L. part. adj. *reducens* bringing back, leading back; N.L. part. adj. *glycinireducens* glycine-reducing).
Psychroacetigena (Psy.chro.a.ce'ti.ge'na. Gr. adj. *psychros* cold; N.L. n. *acidum aceticum* acetic acid; L. suff. -*genus -a -um* (from L. v. *gigno*) producing acetate; N.L. n. *Psychroacetigena* producing acetate under psychrophilic conditions). Psychroacetigena formicireducens (for.mi.ci.re.du'cens. *N.L. n. acidum formicum* formic acid; L. part. adj. *reducens* bringing back, leading back; N.L. part. adj. *formicireducens* formate-reducing).

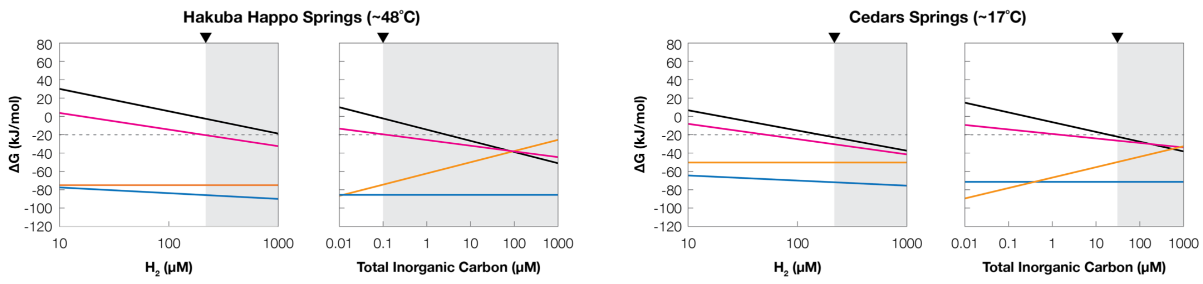


**Figure S1.** Thermodynamics of $H_2$-oxidizing reduction of $CO_2$ (black), formate (pink), and glycine (blue), and formate disproportionation (orange) with changing $H_2$ and total inorganic carbon (TIC) concentrations. The maximum observed $H_2$ concentration in Hakuba Happo hot springs (664 µM) is marked (triangles) in the $H_2$-based plots and the TIC concentrations observed in Hakuba (<0.1 µM) and The Cedars GPS1 (35 µM) are indicated correspondingly.

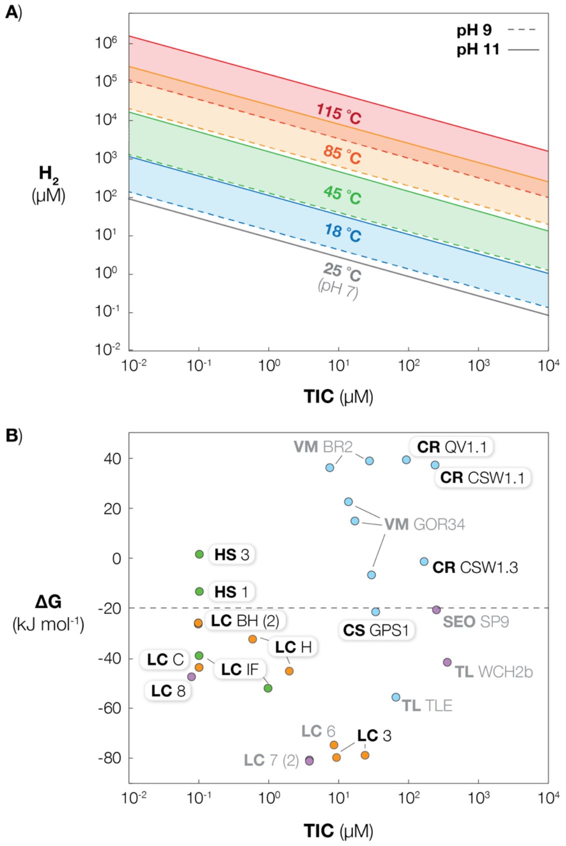


**Figure S2.** Thermodynamics of $H_2$-oxidizing $CO_2$-reducing homoacetogenesis under differing pH, temperature, $H_2$, and total inorganic carbon (TIC) concentrations. (A) For temperatures 18 (blue), 45 (green), 85 (orange), and 115 (red) °C, the $H_2$ and TIC concentrations at which $H_2/CO_2$ homoacetogenesis has a Gibbs free energy yield (ΔG) of -10 kJ $mol^{-1}$ is shown for *p*H of 9 (dotted line) and 11 (solid line) (atmospheric pressure of 1 atm). For reference, the same is shown for 25 °C at *p*H 7 (gray solid line). (B) The ΔG of $H_2/CO_2$ homoacetogenesis in various serpentinite-hosted systems (Hakuba Happo hot springs - HS; The Cedars springs - CS; Lost City - LC; Voltri Massif - VM; Coast Range Ophiolite Microbiological Observatory - CR; Santa Elena ophiolite - SEO; Table Lands - TLE) are shown based on reported environmental conditions for individual sampling locations. Each data point is colored based on temperature: psychrophilic (blue), mesophilic (purple), thermophilic (green), and hyperthermophilic (orange). Samples with associated acetate measurements are labelled black, and those that have >2 µM acetate are circled. For samples with no reported acetate concentrations (gray), the average of reported concentrations was used (8.57 µM Acetate). For The Cedars spring sample, no $H_2$ concentration has been reported, so the highest on-land serpentinite-hosted system $H_2$ concentration was used (664 µM $H_2$ from Hakuba Happo #1). Thermodynamic calculations were performed using $\Delta G°_f$ and $\Delta H°_f$ values at 298 K values and temperature adjustment through the Gibbs-Helmholtz equation.

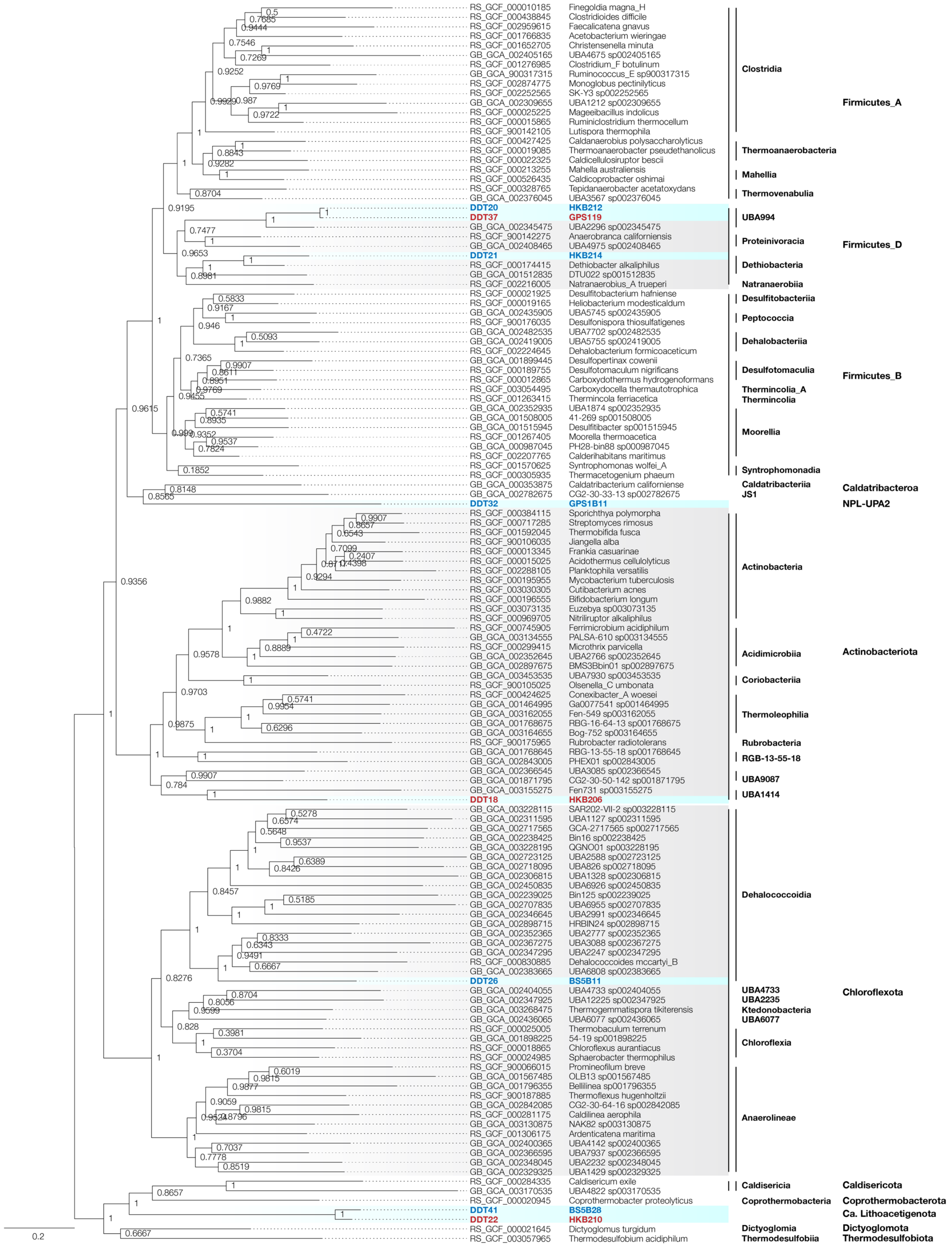


**Figure S3.** Ribosomal protein tree for high-quality MAGs. Universally conserved ribosomal proteins were collected from each genome, aligned with MAFFT v7.394 (Katoh et al., Nucleic Acids Res 30(14), 3059-3066, 2002), trimmed with trimAl 1.2rev59 (-gt 0.70) Capella-Gutierrez et al., Bioinformatics 25(15), 1972–1973, 2009), and concatenated. A maximum likelihood tree was calculated using phyML 3.3.20190321 with the LG model and 100 bootstrap iterations (Guindon and Gascuel, Systematic Biology 52(5), 696-704, 2003). GTDBtk-based phylogeny is shown.

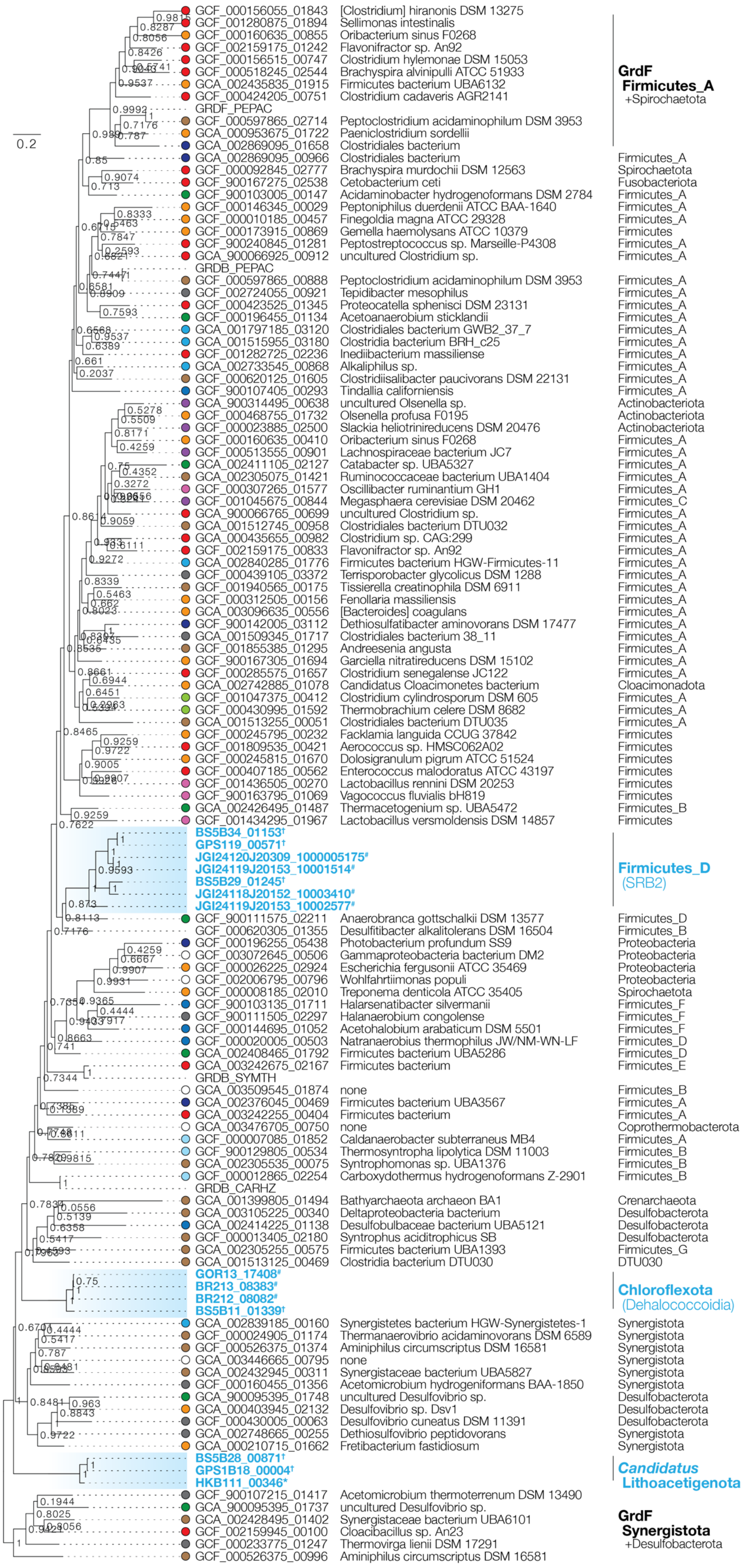


**Figure S4.** Complete phylogenetic tree of glycine reductase subunit GrdB and homologs from Hakuba Happo hot spring*, The Cedars springs†, and other serpentinite-hosted system metagenomes#. COG1978 homologs were collected from the representative species genomes in GTDB, filtered using a GrdB motif conserved across members of phyla known to perform glycine-reducing Stickland reaction (see Fig. 2) and a GrdF motif conserved across sequences that form a distinct cluster around the biochemically characterized *Peptoclostridium acidaminophilum* sarcosine reductase subunit GrdF (see Fig. 2), clustered with 75% amino acid sequence similarity using CD-HIT (-c 0.75), aligned with MAFFT v7.453 (default parameters), and trimmed through trimAl v1.2rev59 (-gt 0.9). A maximum likelihood tree was constructed through PhyML v3.3.20190909 using the LG model and 100 bootstrap replicates (-b 100 -d aa -m LG -v e --leave_duplicates). Bootstrap values were recalculated using BOOSTER. For each sequence, the original habitat the isolate or MAG was obtained from is shown. Large sequence clusters were grouped (number of representative sequences included are shown). Note that the outgroup is a cluster of uncharacterized *Synergistetes* and *Deltaproteobacteria* sequences that was inferred to function as GrdF given that it shares the motif found in the *Firmicutes* GrdF.

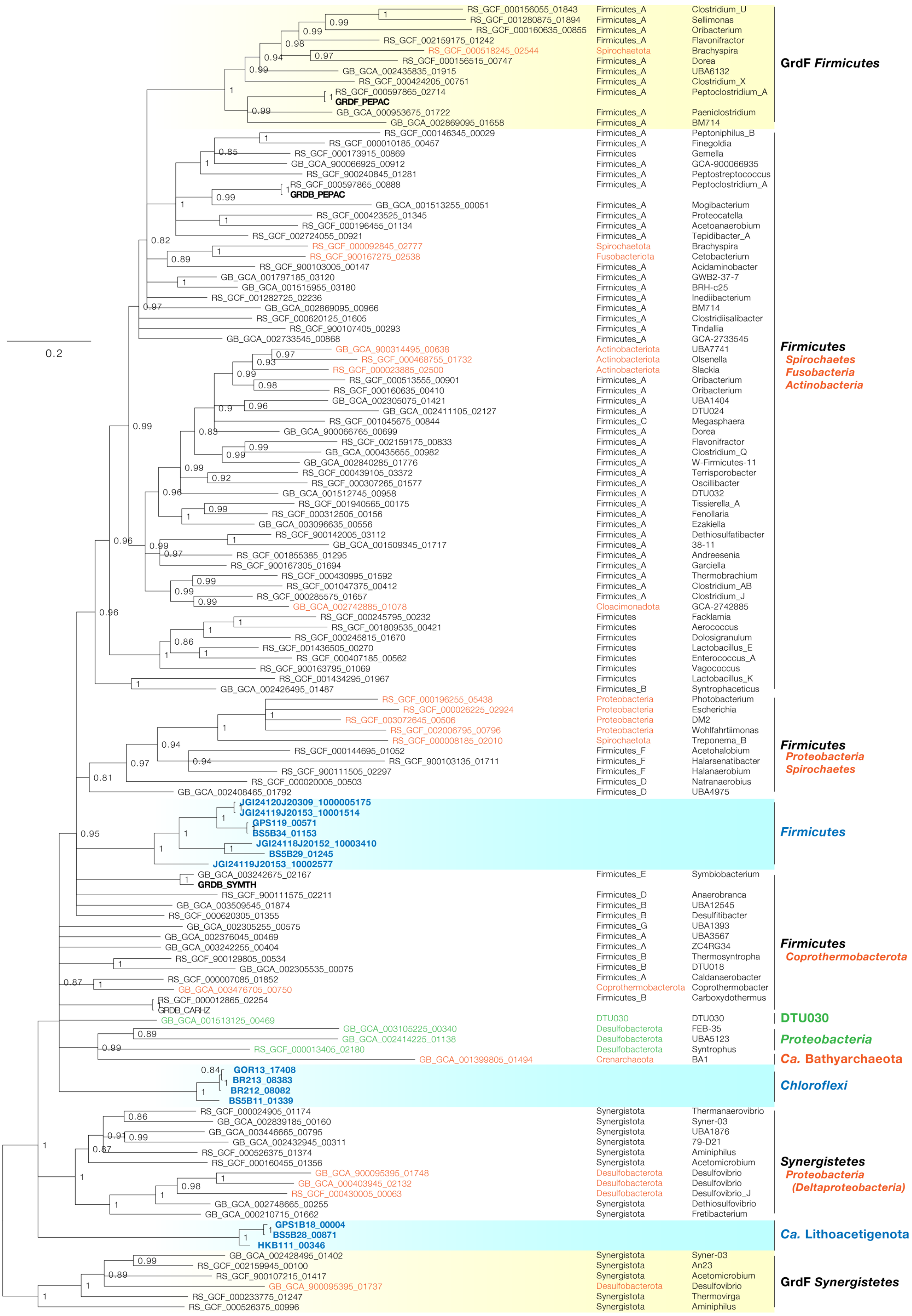

**Figure S5.** Phylogenetic tree of GrdB constructed from bayesian inference. The consensus tree was calculated based on three chains (length of 10000) with a burn-in of 1000 using Phylobayes v1.8. Branches with posterior probabilities less than 0.8 were collapsed. Sequences obtained from serpenitite-hosted sites (blue), likely obtained horizontally (orange), and with unclear phylogenetic positions (green) are shown.

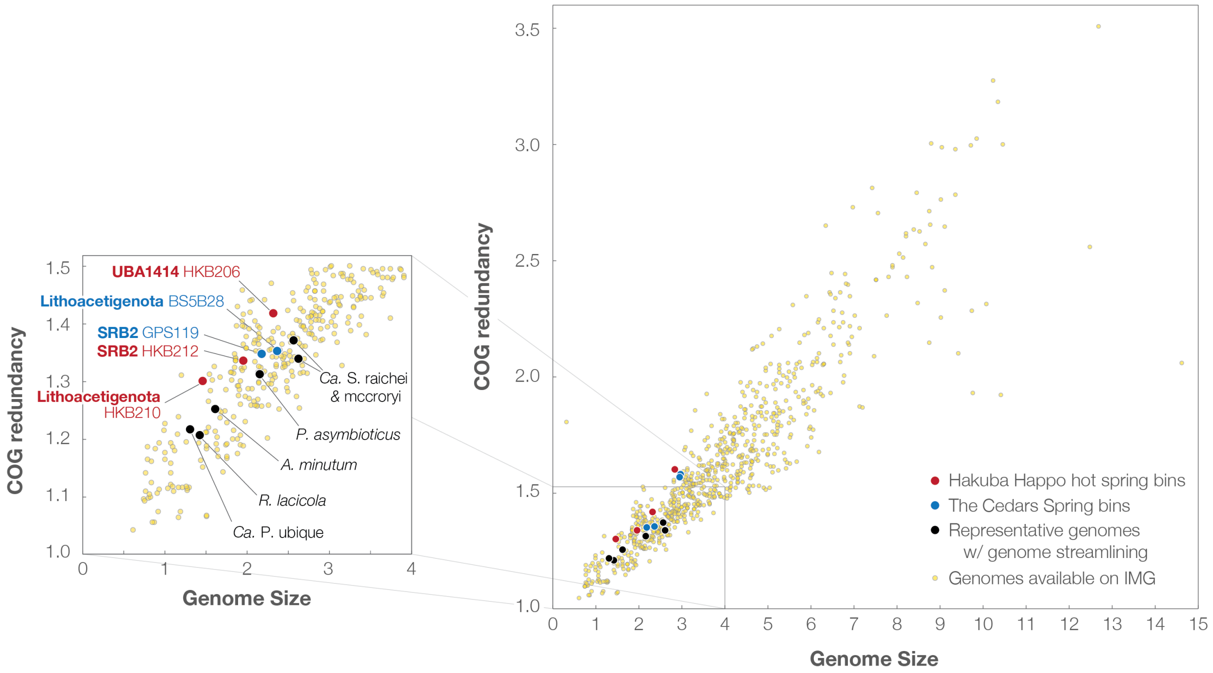


**Figure S6.** Genome streamlining of publically available genomes (Joint Genome Institute Integrated Microbial Genome) and selected high completeness bins from Hakuba Happo hot springs (red) and The Cedars springs (blue) (inset on left). Genomes with known streamlining are marked (black): *Aurantimicrobium minutum*, *Ca.* Pelagibacter ubique, *Polynucleobacter asymbioticus*, *Rhodoluna lacicola*, *Ca.* Serpentinomonas raichei, and *Ca.* S. mccroryi.

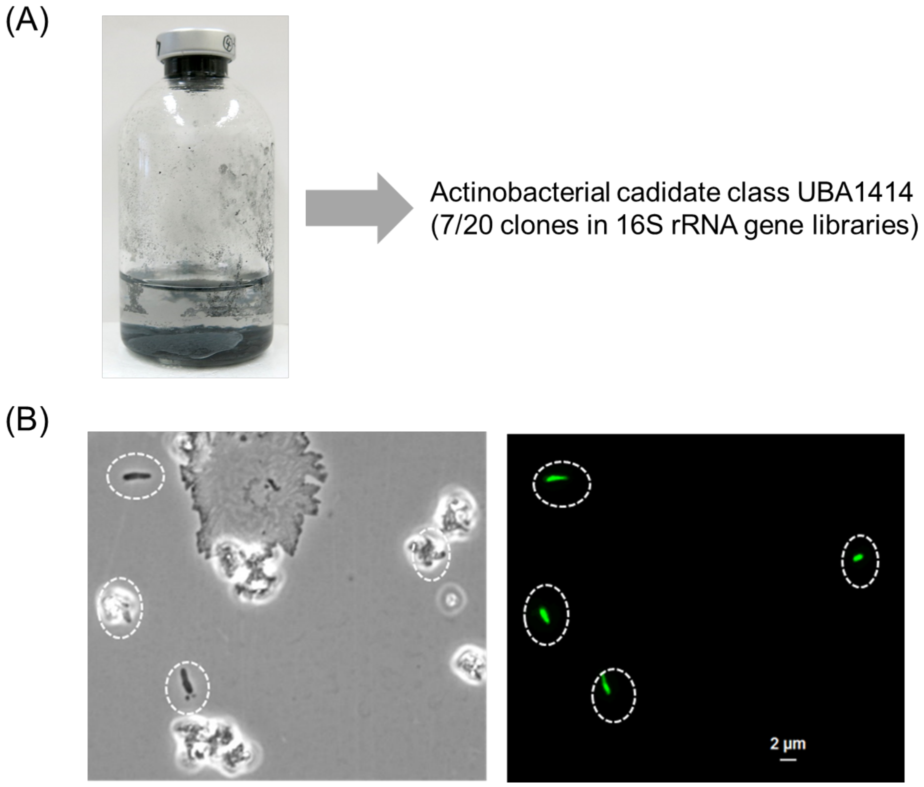


**Figure S7.** Hakuba-derived enrichment culture of *Actinobacteria* UBA1414. (A) Culture medium based on Widdel medium (*p*H 10) with an $N_2$-$CO_2$ (80:20, vol/vol) headspace was supplemented with 0.01 g $l^{-1}$ yeast extract and 25 g $l^{-1}$ elemental iron granules. Hakuba hot spring water 100 mL was passed though a membrane filter and the filter was submerged in the culture medium. After a 4 month incubation at 40 °C, 1 mL of the culture was used for DNA extraction, PCR amplification, and clone library construction (about 600 bp of 16S rRNA gene). UBA1414-derived 16S rRNA gene fragments, which shared high sequence identity (>99%) to bin HKB206, comprised 7 out of 20 clones. The remaining 13 clones consisted of obligately aerobic *Methylobacterium*- and *Pseudomonas*-related sequences that may be contaminants, but further investigation is required. (B) Micrographs: phase-contrast (left) and SYBR-Green-I-stained microbial cells (green, right); scale bar, 2 μm.

**Table S1.** Phylogeny and quality of bins from Hakuba Happo hot springs and The Cedars springs (GPS1 and BS5). Phylogeny was defined by GTDBtk (g) or comparing GTDBtk-defined phylogeny with EMBL (e) or SILVA (s). GTDBtk annotations with RED valu than 0.5 were not considered and phylogeny was checked by constructing a concatenated ribosomal protein tree (see Supplement). * Low quality bins that were only used for comparative purposes (e.g., whether a function found in a high-quality bin from HKE present in HKB2 with >99% similarity)

| Information | | | | Phylogeny | | | | Genome statistics | | |
|---|---|---|---|---|---|---|---|---|---|---|
| **Habitat** | **Sample** | **Bin** | **Prefix** | **Phylogeny (phylum)** | **Phylogeny (lowest level)** | **based on GTDBtk** | **GTDBtk RED value** | **Genome size (Mb)** | **Contigs (#)** | **Complete ness (%)** |
| Hakuba | HKB702 | HKB206 | DDT18 | *Actinobacteria* | UBA1414 (g) | d__Bacteria;p__Actinobacteriota;c__UBA1414;o__;f__;g__;s__ | 0.502308 | 1.97 | 346 | 85.5 |
| Hakuba | HKB701 | HKB109 | DDT19 | *Firmicutes* | *Syntrophomonadaceae* (e) | d__Bacteria;p__Firmicutes_D;c__Dethiobacteria;o__Dethiobacterales;f__Dethiobacteraceae;g__;s__ | 0.850189 | 2.98 | 562 | 73.2 |
| Hakuba | HKB702 | HKB212 | DDT20 | *Firmicutes* | SRB2 (s) | d__Bacteria;p__Firmicutes_D;c__UBA994;o__UBA994;f__UBA994;g__UBA994;s__ | 0.937622 | 1.75 | 209 | 89.4 |
| Hakuba | HKB702 | HKB214 | DDT21 | *Firmicutes* | *Syntrophomonadaceae* (e) | d__Bacteria;p__Firmicutes_D;c__Dethiobacteria;o__Dethiobacterales;f__;g__;s__ | 0.686205 | 2.67 | 112 | 89.4 |
| Hakuba | HKB702 | HKB210 | DDT22 | Ca. Lithoacetigenota (novel) | - | *d__Bacteria;p__Coprothermobacterota;c__Coprothermobacteria;o__;f__;g__;s__* | 0.417028 | 1.30 | 223 | 88.8 |
| Hakuba | HKB701 | HKB111 | DDT23 | Ca. Lithoacetigenota (novel) | - | *d__Bacteria;p__Coprothermobacterota;c__Coprothermobacteria;o__;f__;g__;s__* | 0.421002 | 1.24 | 204 | 83.6 |
| Cedars | GPS1 2011 | GPS105* | DDT24 | *Chloroflexi* | *Dehalococcoidia* (s) | d__Bacteria;p__Chloroflexota;c__Dehalococcoidia;o__SZUA-161;f__;g__;s__ | 0.632993 | 0.88 | 233 | 62 |
| Cedars | BS5 2011 | BS517* | DDT25 | *Chloroflexi* | *Dehalococcoidia* (s) | d__Bacteria;p__Chloroflexota;c__Dehalococcoidia;o__SZUA-161;f__;g__;s__ | 0.629093 | 1.31 | 304 | 59.7 |
| Cedars | BS5 2012 | BS5B11 | DDT26 | *Chloroflexi* | *Dehalococcoidia* (s) | d__Bacteria;p__Chloroflexota;c__Dehalococcoidia;o__SZUA-161;f__;g__;s__ | 0.628746 | 2.51 | 486 | 85.1 |
| Cedars | BS5 2011 | BS503 | DDT27 | *Chloroflexi* | *Dehalococcoidia* (s) | d__Bacteria;p__Chloroflexota;c__Dehalococcoidia;o__SZUA-161;f__;g__;s__ | 0.626397 | 1.52 | 342 | 74.1 |
| Cedars | GPS1 2012 | GPS1B04* | DDT28 | *Chloroflexi* | *Dehalococcoidia* (s) | d__Bacteria;p__Chloroflexota;c__Dehalococcoidia;o__SZUA-161;f__;g__;s__ | 0.634978 | 1.13 | 298 | 59.1 |
| Cedars | GPS1 2011 | GPS109* | DDT29 | *Firmicutes* | *Syntrophomonadaceae* (e) | d__Bacteria;p__Firmicutes_D;c__Dethiobacteria;o__Dethiobacterales;f__Dethiobacteraceae;g__;s__ | 0.8487 | 2.70 | 688 | 85.2 |
| Cedars | GPS1 2012 | GPS1B09 | DDT30 | *Firmicutes* | *Syntrophomonadaceae* (e) | d__Bacteria;p__Firmicutes_D;c__Dethiobacteria;o__Dethiobacterales;f__Dethiobacteraceae;g__;s__ | 0.847308 | 2.32 | 503 | 86.9 |
| Cedars | BS5 2012 | BS5B29 | DDT34 | *Firmicutes* | SRB2 (s) | d__Bacteria;p__Firmicutes_D;c__UBA994;o__UBA994;f__UBA994;g__;s__ | 0.828123 | 2.60 | 521 | 91.5 |
| Cedars | GPS1 2011 | GPS123 | DDT35 | *Firmicutes* | SRB2 (s) | d__Bacteria;p__Firmicutes_D;c__UBA994;o__UBA994;f__UBA994;g__;s__ | 0.824263 | 1.51 | 279 | 92.2 |
| Cedars | BS5 2011 | BS524 | DDT36 | *Firmicutes* | SRB2 (s) | d__Bacteria;p__Firmicutes_D;c__UBA994;o__UBA994;f__UBA994;g__;s__ | 0.820371 | 1.74 | 227 | 88.1 |
| Cedars | GPS1 2011 | GPS119 | DDT37 | *Firmicutes* | SRB2 (s) | d__Bacteria;p__Firmicutes_D;c__UBA994;o__UBA994;f__UBA994;g__UBA994;s__ | 0.937791 | 1.95 | 283 | 89.41 |
| Cedars | BS5 2011 | BS530 | DDT38 | *Firmicutes* | SRB2 (s) | d__Bacteria;p__Firmicutes_D;c__UBA994;o__UBA994;f__UBA994;g__UBA994;s__ | 0.941416 | 1.68 | 384 | 86.4 |
| Cedars | BS5 2012 | BS5B34 | DDT39 | *Firmicutes* | SRB2 (s) | d__Bacteria;p__Firmicutes_D;c__UBA994;o__UBA994;f__UBA994;g__UBA994;s__ | 0.934183 | 1.69 | 329 | 88.5 |
| Cedars | BS5 2011 | BS529* | DDT31 | NPL-UPA2 | - | *d__Bacteria;p__Ratteibacteria;c__UBA8468;o__;f__;g__;s__* | 0.46067 | 2.02 | 481 | 74.4 |
| Cedars | GPS1 2012 | GPS1B11 | DDT32 | NPL-UPA2 | - | *d__Bacteria;p__;c__;o__;f__;g__;s__* | *0.381487* | 2.47 | 583 | 87 |
| Cedars | GPS1 2011 | GPS112 | DDT33 | NPL-UPA2 | - | *d__Bacteria;p__;c__;o__;f__;g__;s__* | *0.382715* | 1.81 | 401 | 85.3 |
| Cedars | GPS1 2012 | GPS1B18* | DDT40 | Ca. Lithoacetigenota (novel) | - | *d__Bacteria;p__Coprothermobacterota;c__Coprothermobacteria;o__;f__;g__;s__* | *0.417496* | 2.06 | 596 | 79.31 |
| Cedars | BS5 2012 | BS5B28 | DDT41 | Ca. Lithoacetigenota (novel) | - | *d__Bacteria;p__Coprothermobacterota;c__Coprothermobacteria;o__;f__;g__;s__* | *0.42171* | 1.75 | 278 | 94.8 |
| Cedars | BS5 2011 | BS525 | DDT42 | Ca. Lithoacetigenota (novel) | - | *d__Bacteria;p__Coprothermobacterota;c__Coprothermobacteria;o__;f__;g__;s__* | *0.417261* | 2.12 | 427 | 89.7 |

**Table S2. Distribution of acetogenesis-related pathways among bins recovered from Hakuba Happo hot springs and The Cedars springs.** Low quality bins are grayed out. * Missing one non-substrate-binding subunit of a multi-subunit protein complex.

| Bin | Prefix | Phylum | Specific | Hydrogenase | | | | | Formate dehydrogenase | | | | | THF pathway | | | | | CO deh. | Bacterial CODH/ACS | | | | | Archaeal CODH/ACS | | | | | Hybrid CODH/ACS | | | | Glycine reductase | | Electron transport | | | | | | | | | |
|---|---|---|---|---|---|---|---|---|---|---|---|---|---|---|---|---|---|---|---|---|---|---|---|---|---|---|---|---|---|---|---|---|---|---|---|---|---|---|---|---|---|---|---|---|---|
| | | | | hoxEFUHY | hoxEFUHYP | hydABC | hydABCD | hndABCD | fdhA-hylBC | fdhA-hylABC | fdhA | fdpAB | fdnGH-hybAB | fhs | folD | fchA | metV | metF | CooS | acsA | acsB | acsC | acsD | acsE | cdhA | cdhB | cdhC | cdhD | cdhE | acsABCDE | cdhABCD-acsCE | cdhABC-acsCDE | cdhBC-acsACDE | grdABCDE | trxAB | rnfA | rnfB | rnfC | rnfD | rnfE | rnfG | nfnAB | hdrA | hdrB | hdrC |
| GPS109 | DDT29 | *Firmicutes* | *Syntrophomonadaceae (e)* | | | | + | | | | + | | | + | + | | + | + | | + | + | + | + | + | + | + | + | + | | + | + | + | | | | + | + | + | + | + | + | | | | |
| GPS1B09 | DDT30 | *Firmicutes* | *Syntrophomonadaceae (e)* | | | + | + | | | | | | | + | + | | + | + | + | + | | + | + | + | + | + | + | | | | | + | | | | + | + | + | + | + | + | | | | |
| HKB109 | DDT19 | *Firmicutes* | *Syntrophomonadaceae (e)* | | | + | | | | | | | | + | + | | + | + | | + | + | + | + | + | + | + | + | + | | + | + | + | | | | + | + | + | + | + | + | + | + | | |
| HKB214 | DDT21 | *Firmicutes* | *Syntrophomonadaceae (e)* | | | + | + | | | + | | + | | + | + | | + | + | + | + | + | + | + | + | + | + | + | + | | + | + | + | | | | + | + | + | + | + | + | | + | + | + |
| GPS1B11 | DDT32 | NPL-UPA2 | - | | | + | | | + | | | | | + | + | | + | + | + | + | + | + | + | + | + | + | + | + | | + | + | + | | | | + | + | + | + | + | + | | + | | |
| GPS112 | DDT33 | NPL-UPA2 | - | | | | | | | | | | | + | + | | + | + | | | | + | + | + | + | + | + | | | | | | | | | + | + | + | + | + | + | | + | | |
| BS529 | DDT31 | NPL-UPA2 | - | | | + | | | | | | | | | + | | + | + | | | | + | | + | | | | | | | | | | | | + | | + | + | + | + | | + | + | |
| HKB206 | DDT18 | *Actinobacteria* | UBA1414 (g) | | + | | | | + | | | | | + | + | | + | + | + | + | | + | + | + | | + | + | | | | | | + | | | + | + | | + | + | + | + | + | + | + |
| BS525 | DDT42 | Ca. Lithoacetigenota (novel) | - | | + | | | | | | | | | + | + | | + | + | | | | + | + | + | + | + | + | | | | | + | | + | + | + | + | + | + | + | + | | + | + | |
| BS5B28 | DDT41 | Ca. Lithoacetigenota (novel) | - | | + | | | | | | | | | + | + | | + | + | | | | + | + | + | + | + | + | | | | | + | | + | + | + | + | + | + | + | + | | + | + | |
| GPS1B18 | DDT40 | Ca. Lithoacetigenota (novel) | - | | + | | | | | | | | | + | + | | + | | | | | + | + | + | + | + | + | | | | | + | | + | + | + | + | + | | + | + | | + | + | |
| GPS105 | DDT24 | *Chloroflexi* | *Dehalococcoidia (s)* | | | + | | | | + | | | | | | | + | + | | + | | | + | + | + | + | | + | | | | | | | | + | | + | | | | + | + | | |
| GPS1B04 | DDT28 | *Chloroflexi* | *Dehalococcoidia (s)* | + | | + | | | | | + | | | | | + | + | + | + | | | | | | | | | | | | | | | +* | + | + | | + | | | + | | + | | |
| BS503 | DDT27 | *Chloroflexi* | *Dehalococcoidia (s)* | | | + | | | | | + | | | + | + | | | + | | + | | | + | + | | | | + | | | | | | | | + | + | + | + | | + | | + | + | |
| BS517 | DDT25 | *Chloroflexi* | *Dehalococcoidia (s)* | | | | | | + | | | | | + | | | + | + | + | | | | | | | | | | | | | | | | | | | + | | | | + | + | | |
| BS5B11 | DDT26 | *Chloroflexi* | *Dehalococcoidia (s)* | | | + | | | | + | | | | + | + | + | + | + | | | | + | + | + | | + | | + | | | | | | +* | + | + | + | + | + | | + | | + | + | |
| BS530 | DDT38 | *Firmicutes* | SRB2 (s) | | | + | + | | + | + | | | | + | | + | | | | | | | | | | | | | | | | | | | | + | + | | + | + | + | | + | | |
| GPS119 | DDT37 | *Firmicutes* | SRB2 (s) | | | + | + | | | | | | | + | | + | | | | | | | | | | | | | | | | | | + | + | + | + | + | + | + | + | | | | |
| HKB212 | DDT20 | *Firmicutes* | SRB2 (s) | | | + | + | | | | | | | + | + | + | | | | | | | | | | | | | | | | | | + | + | + | + | + | + | + | + | | | | |
| BS5B34 | DDT39 | *Firmicutes* | SRB2 (s) | | | + | | | | | | | | + | | + | | | | | | | | | | | | | | | | | | + | + | | + | | + | | | | | | |
| BS5B29 | DDT34 | *Firmicutes* | SRB2 (s) | | | + | + | | + | + | | | | + | + | + | | | | | | | | | | | | | | | | | | + | + | + | | + | + | + | + | | + | | |
| GPS123 | DDT35 | *Firmicutes* | SRB2 (s) | | | + | | | | | | | | + | + | + | | | | | | | | | | | | | | | | | | | | + | + | | + | + | + | | | | |
| BS524 | DDT36 | *Firmicutes* | SRB2 (s) | | | + | | + | | | | | | + | + | + | | | | | | | | | | | | | | | | | | | | + | + | + | + | + | + | | | | |
| HKB111 | DDT23 | Ca. Lithoacetigenota (novel) | - | | +* | | | | | | | | | | | | | | | | | | | + | | | | | | | | | | + | + | + | + | + | + | + | + | | + | | |
| HKB210 | DDT22 | Ca. Lithoacetigenota (novel) | - | | + | | | | | | | | | + | | | | | | | | | + | + | | | | | | | | | | + | + | + | + | + | + | + | + | | | | |

**Table S3. Locus tags for acetogenesis-related genes in select Hakuba Happo hot spring and The Cedars springs bins.**

| | | Ca. Lithoacetigenota | | UBA1414 | NPL-UPA2 | *Syntrophomonadaceae* | *Syntrophomonadaceae* | *Syntrophomonadaceae* | *Dehalococcoidia* | SRB2 | SRB2 |
|---|---|---|---|---|---|---|---|---|---|---|---|
| | | HKB210 | BS5B28 | HKB206 | GPS1B11 | HKB214 | GPS1B09 | HKB109 | BS5B11 | HKB212 | GPS119 |
| **Glycine reductase** | GrdE-TrxA | DDT22_00297 | DDT41_00874 | - | - | - | - | - | - | - | - |
| | GrdE | *DDT22_00297* | *DDT41_00874* | - | - | - | - | - | DDT26_01340 | DDT20_01364 | DDT37_00574 |
| | GrdA1 | DDT22_00296 | DDT41_00873 | - | - | - | - | - | DDT26_01344 | DDT20_01365 | DDT37_00573 |
| | GrdA2 | DDT22_00295 | DDT41_00872 | - | - | - | - | - | DDT26_01343 | DDT20_01366 | DDT37_00572 |
| | GrdB | DDT22_00294 | DDT41_00871 | - | - | - | - | - | DDT26_01339 | DDT20_01367 | DDT37_00571 |
| | GrdC | DDT22_00292 | DDT41_00869 | - | - | - | - | - | - | *DDT20_01369* | *DDT37_00570* |
| | GrdD | DDT22_00291 | DDT41_00868 | - | - | - | - | - | - | *DDT20_01369* | *DDT37_00570* |
| | GrdCD | - | - | - | - | - | - | - | - | DDT20_01369 | DDT37_00570 |
| **Thioredoxin** | TrxA | *DDT22_00297* | *DDT41_00874* | - | - | - | - | - | DDT26_00572 | DDT20_01363 | DDT37_00575 |
| | TrxB | DDT22_00328 | DDT41_00629 | - | - | - | - | - | DDT26_00593 | DDT20_01362 | DDT37_00576 |
| **Sec biosynthesis** | SelA | DDT22_00171 | DDT41_00605 | DDT18_01429 | - | DDT21_01805 | DDT30_01458 | - | DDT26_01079 | DDT20_01332 | DDT37_00583 |
| | SelB | DDT22_00172 | DDT41_00604 | DDT18_01428 | - | DDT21_01806 | DDT30_01459 | - | DDT26_00526 | DDT20_01331 | DDT37_00582 |
| **NiFe hydrogenase** | HoxY | DDT22_00350 | DDT41_00425 | DDT18_00660 | - | - | - | - | DDT26_02429 | - | - |
| | HoxH | DDT22_00351 | DDT41_00426 | DDT18_00659 | - | - | - | - | - | - | - |
| | HoxU | DDT22_00349 | DDT41_00424 | DDT18_00661 | - | - | - | - | DDT26_01928 | - | - |
| | HoxF | DDT22_00348 | DDT41_00423 | DDT18_00626 | - | - | - | - | DDT26_00808 | - | - |
| | HoxE | DDT22_00347 | DDT41_00422 | DDT18_00627 | - | - | - | - | DDT26_00809 | - | - |
| | HoxP | DDT22_00346 | DDT41_00421 | DDT18_00662 | - | - | - | - | - | - | - |
| **FeFe hydrogenase** | HydA | - | - | - | - | DDT21_01978 | DDT30_01055 | - | DDT26_01129 | - | - |
| | HydB | - | - | - | - | DDT21_01977 | DDT30_01054 | - | DDT26_01128 | - | - |
| | HydC | - | - | - | - | DDT21_01976 | DDT30_01053 | - | DDT26_01127 | - | - |
| | HydA | - | - | - | - | - | - | - | - | DDT20_00909 | DDT37_00389 |
| | HydB | - | - | - | - | - | - | - | - | DDT20_00908 | DDT37_00388 |
| | HydC | - | - | - | - | - | - | - | - | DDT20_00907 | DDT37_00387 |
| | HydD | - | - | - | - | - | - | - | - | DDT20_00906 | DDT37_00386 |
| | HydA | - | - | - | - | - | - | - | - | - | DDT37_00655 |
| | HydB | - | - | - | - | - | - | - | - | - | DDT37_00654 |
| | HydC | - | - | - | - | - | - | - | - | - | DDT37_00653 |
| | HydD | - | - | - | - | - | - | - | - | - | DDT37_00652 |
| | HndA | - | - | - | - | DDT21_01026 | DDT30_00515 | DDT19_00831 | - | - | - |
| | CckA | | | | | DDT21_01027 | DDT30_00514 | DDT19_00832 | - | | |
| | HndB | - | - | - | - | DDT21_01028 | DDT30_00513 | DDT19_00833 | - | - | - |
| | HndC | - | - | - | - | DDT21_01029 | DDT30_00512 | DDT19_00834 | - | - | - |
| | HndD | - | - | - | - | DDT21_01030 | DDT30_00511 | DDT19_00835 | - | - | - |
| **Formate dehdyrogenase** | FdhA | - | - | DDT18_01054 | DDT32_00315 | DDT21_02578 | - | - | DDT26_01503 | - | - |
| | (FdhA2) | - | - | - | - | DDT21_02579 | - | - | DDT26_01504 | - | - |
| | HylB | - | - | DDT18_01056 | DDT32_00311 | DDT21_02576 | - | - | DDT26_01501 | - | - |
| | HylC | - | - | DDT18_01055 | DDT32_00312 | DDT21_02575 | - | - | DDT26_01502 | - | - |
| | MetV | - | - | DDT18_01057 | DDT32_00310 | - | - | - | - | - | - |
| | MetF | - | - | - | DDT32_00309 | - | - | - | - | - | - |
| | FdhA | - | - | - | - | DDT21_01601 | - | - | - | - | - |
| | (FdhA2) | - | - | - | - | - | - | - | - | - | - |
| | FdpB | - | - | - | - | DDT21_01599 | - | - | - | - | - |
| | FdhA | - | - | - | - | - | - | - | - | - | - |
| | (FdhA2) | - | - | - | - | - | - | - | - | - | - |
| | FdpAB | - | - | - | - | DDT21_01162 | - | - | - | - | - |
| | FdnG1 | - | - | - | - | - | - | - | - | - | - |
| | FdnG2 | - | - | - | - | - | - | - | - | - | - |
| | FdnH/HybA | - | - | - | - | - | - | - | - | - | - |
| | HybA | - | - | - | - | - | - | - | - | - | - |
| | HybB | - | - | - | - | - | - | - | - | - | - |
| **C1 metabolism** | FchA | - | - | - | - | - | - | - | DDT26_02662 | DDT20_01400 | DDT37_00119 |
| | Fhs | DDT22_01344 | DDT41_01181 | DDT18_01669 | DDT32_00091 | DDT21_00652 | DDT30_01889 | DDT19_01668 | DDT26_02308 | DDT20_01401 | DDT37_00120 |
| | FolD | - | DDT41_01485 | DDT18_01668 | DDT32_00071 | DDT21_01161 | DDT30_01019 | DDT19_01973 | DDT26_02307 | DDT20_00767 | - |
| | MetV | - | DDT41_01411 | DDT18_00798 | DDT32_00310 | DDT21_00074 | DDT30_00161 | DDT19_00973 | DDT26_01099 | - | - |
| | MetF | - | DDT41_01410 | DDT18_00797 | DDT32_00309 | DDT21_00075 | DDT30_00160 | DDT19_00974 | DDT26_01100 | - | - |
| | MtmB | - | - | DDT18_01085 | - | - | - | - | - | - | - |
| | MtmC | - | - | DDT18_01084 | - | - | - | - | - | - | - |
| **Acetyl-CoA synthase / CO dehydrogenase** | CooS | - | - | DDT18_01860(?) | DDT32_02123 | DDT21_01084 | DDT30_01952 | - | - | - | - |
| | AcsA | - | - | - | DDT32_00844 | DDT21_00067 | DDT30_01473 | DDT19_01000 | - | - | - |
| | AcsB | - | - | - | DDT32_00845 | DDT21_00068 | - | DDT19_02607 | - | - | - |
| | AcsC | - | - | - | DDT32_00795 | DDT21_00069 | DDT30_00165 | DDT19_00968 | - | - | - |
| | AcsD | - | - | - | DDT32_00843 | DDT21_00071 | DDT30_00164 | DDT19_00970 | DDT26_02597 | - | - |
| | AcsE | - | - | - | DDT32_00794 | DDT21_00072 | DDT30_00163 | DDT19_00971 | DDT26_02596 | - | - |
| | CdhA | - | DDT41_00748 | - | DDT32_00095 | DDT21_00063 | DDT30_00615 | DDT19_02581 | - | - | - |
| | CdhB | - | DDT41_00750 | - | DDT32_00094 | DDT21_00064 | DDT30_00616 | DDT19_02580 | - | - | - |
| | CdhC | - | DDT41_00751 | - | DDT32_00093 | DDT21_00065 | DDT30_00617 | - | - | - | - |
| | AcsC | - | DDT41_00752 | - | DDT32_00092 | DDT21_00066 | DDT30_00618 | - | - | - | - |
| | AcsD | - | DDT41_01017 | - | DDT32_00096 | DDT21_00062 | DDT30_00614 | - | - | - | - |
| | AcsE | - | DDT41_01016 | - | DDT32_00098 | DDT21_00061 | DDT30_00613 | - | - | - | - |
| | AcsA | - | - | DDT18_00628 | - | - | - | - | - | - | - |
| | CdhB | - | - | DDT18_01643 | - | - | - | - | - | - | - |
| | CdhC | - | - | DDT18_01644 | - | - | - | - | - | - | - |
| | AcsC1 | - | - | DDT18_01645 | - | - | - | - | - | - | - |
| | AcsC2 | - | - | DDT18_01462 | - | - | - | - | - | - | - |
| | AcsD1 | - | - | DDT18_01460 | - | - | - | - | - | - | - |
| | AcsD2 | - | - | DDT18_01461 | - | - | - | - | - | - | - |
| | AcsE | - | - | DDT18_00130 | - | - | - | - | - | - | - |
| **Substrate-level phosphorylation** | Pta | DDT23_18: 12235-127161 | DDT41_00043 | DDT18_01737 | - | - | - | - | - | DDT20_01393 | DDT37_00439 |
| | Ack | DDT22_00702 | DDT41_00042 | - | - | - | - | - | - | DDT20_01392 | DDT37_00438 |
| | Ack | - | DDT41_00044 | - | - | - | - | - | - | DDT20_01394 | DDT37_00441 |
| | Acs1 | - | DDT41_01004 | DDT18_01738* | DDT32_00571 | DDT21_01032 | DDT30_00052 | - | DDT26_01691 | - | - |
| | Acs2 | - | - | - | DDT32_00005 | DDT21_01574 | - | - | DDT26_01010 | - | - |
| | ActP | - | - | DDT18_01988 | - | DDT21_00671 | - | - | - | - | - |
| | (split) | - | - | DDT18_01987 | - | - | - | - | - | - | - |
| **Antiporters** | MrpB | - | - | - | - | DDT21_00144 | DDT30_00118 | DDT19_01170 | DDT26_01065 | DDT20_00034 | DDT37_00374 |
| | MrpC | - | - | - | - | DDT21_00145 | DDT30_00117 | DDT19_01171 | DDT26_01066 | DDT20_00033 | DDT37_00373 |
| | MrpD | - | - | - | - | DDT21_00146 | DDT30_00116 | DDT19_01172 | DDT26_01067 | DDT20_00032 | DDT37_00372 |
| | MrpE | - | - | - | - | DDT21_00147 | DDT30_00115 | DDT19_01173 | DDT26_01068 | DDT20_00031 | DDT37_00371 |
| | MrpF | - | - | - | - | DDT21_00148 | DDT30_00114 | DDT19_01174 | DDT26_01069 | DDT20_00030 | DDT37_00370 |
| | MrpG | - | - | - | - | DDT21_00149 | DDT30_00113 | DDT19_01175 | DDT26_01070 | DDT20_00029 | DDT37_00369 |
| | MrpA/D | - | - | - | - | DDT21_00150 | DDT30_00112 | DDT19_01176 | DDT26_01071 | DDT20_00028 | DDT37_00368 |
| | MrpA/D | - | - | - | - | DDT21_00151 | DDT30_00111 | DDT19_01177 | - | - | - |
| | MrpA/D | - | - | - | - | DDT21_00152 | DDT30_00110 | DDT19_01178 | - | - | - |
| | MnhA | DDT22_00283 | DDT41_00383 | - | - | DDT21_00151 | - | - | - | - | - |
| | MnhA? | DDT22_00284 | - | - | - | - | - | - | - | - | - |
| | MnhB | DDT22_00285 | DDT41_00386 | - | - | DDT21_00148 | - | DDT19_01174 | - | - | - |
| | MnhC | DDT22_00286 | DDT41_00385 | - | - | DDT21_00149 | - | DDT19_01175 | - | - | - |
| | MnhD | DDT22_00287 | DDT41_00384 | - | - | DDT21_00150 | - | DDT19_01176 | - | - | - |
| | MnhE | DDT22_00288 | DDT41_00391 | - | - | DDT21_00143 | - | DDT19_01169 | - | - | - |
| | MnhF? | DDT22_00289 | DDT41_00390 | - | - | DDT21_00144 | - | DDT19_01170 | - | - | - |
| | MnhG | DDT22_00290 | DDT41_00389 | - | - | DDT21_00145 | - | DDT19_01171 | - | - | - |
| | NhaD | - | - | - | DDT32_01888 | - | - | - | - | - | - |
| | KefB | - | - | - | - | DDT21_00752 | - | - | - | - | - |
| | KefC | - | - | - | - | DDT21_00753 | - | - | - | - | - |
| | KhtT | - | - | - | - | DDT21_00754 | - | - | - | - | - |
| **Electron transport** | RnfC | DDT22_00128 | DDT41_00161 | - | DDT32_00181 | DDT21_00136 | - | DDT19_01783 | DDT26_00730 | DDT20_00529 | DDT37_00641 |
| | RnfD | DDT22_00127 | DDT41_00160 | DDT18_01397 | DDT32_00180 | DDT21_00137 | DDT30_00125 | DDT19_01782 | DDT26_01058 | DDT20_00528 | DDT37_00640 |
| | RnfG | DDT22_00126 | DDT41_00159 | DDT18_01398 | DDT32_00179 | DDT21_00138 | DDT30_00124 | DDT19_01781 | DDT26_01059 | DDT20_00037 | DDT37_00376 |
| | RnfE | DDT22_00125 | DDT41_00158 | DDT18_01399 | DDT32_00178 | DDT21_00139 | DDT30_00123 | DDT19_01780 | - | DDT20_00527 | DDT37_00639 |
| | RnfA | DDT22_00124 | DDT41_00157 | DDT18_01400 | DDT32_00177 | DDT21_00140 | DDT30_00122 | DDT19_01779 | DDT26_01062 | DDT20_00526 | DDT37_00638 |
| | RnfB | DDT22_00123 | DDT41_00156 | DDT18_01401 | DDT32_00564 | DDT21_00141 | - | DDT19_01778 | DDT26_01063 | DDT20_00525 | DDT37_00637 |
| **Other Energy conservation** | Bcd | - | - | DDT18_01220 | - | - | - | - | - | DDT20_00891 | DDT37_00293 |
| | EtfB | - | - | DDT18_01219 | - | - | - | - | - | DDT20_00890 | DDT37_00292 |
| | EtfA | - | - | DDT18_01218 | - | - | - | - | - | DDT20_00889 | DDT37_00291 |
| **ATP synthase (A-type)** | subunit A | - | DDT41_01328 | DDT18_00494 | DDT32_00513 | DDT21_01718 | DDT30_00226 | DDT19_00643 | DDT26_00842 | DDT20_00644 | DDT37_00801 |
| | subunit B | - | DDT41_01329 | DDT18_00493 | DDT32_00514 | DDT21_01717 | DDT30_00225 | DDT19_00644 | DDT26_00840 | DDT20_00643 | DDT37_00802 |
| | subunit C | DDT22_00643 | - | DDT18_00496 | DDT32_00511 | DDT21_01720 | DDT30_00228 | DDT19_00641 | - | DDT20_00646 | DDT37_00799 |
| | subunit D | - | DDT41_01330 | DDT18_00492 | DDT32_00515 | DDT21_01716 | DDT30_00224 | DDT19_00645 | - | DDT20_00642 | DDT37_00803 |
| | subunit E | DDT22_00644 | DDT41_00678 | DDT18_00497 | DDT32_00510 | DDT21_01721 | DDT30_00227 | DDT19_00640 | - | DDT20_00647 | DDT37_00798 |
| | subunit F | DDT22_00642 | DDT41_01327 | - | - | - | - | - | - | DDT20_00650 | DDT37_00795 |
| | subunit G | - | - | DDT18_00495 | DDT32_00512 | DDT21_01719 | - | DDT19_00642 | - | DDT20_00645 | DDT37_00800 |
| | subunit I | DDT22_00646 | DDT41_00676 | DDT18_00499 | DDT32_00508 | DDT21_01723 | - | - | - | DDT20_00649 | DDT37_00796 |
| | subunit K | DDT22_00645 | DDT41_00677 | DDT18_00498 | DDT32_00509 | DDT21_01722 | DDT30_00230 | DDT19_00639 | DDT26_00848 | DDT20_00648 | DDT37_00797 |
| **ATP synthase (F-type)** | alpha | - | - | - | - | - | - | DDT19_01308 | - | - | - |
| | beta | - | - | - | - | - | - | DDT19_01306 | - | - | - |
| | gamma | - | - | - | - | - | - | DDT19_01307 | - | - | - |
| | delta | - | - | - | - | - | - | DDT19_01309 | - | - | - |
| | epsilon | - | - | - | - | - | - | DDT19_01305 | - | - | - |
| | a | - | - | - | - | - | - | DDT19_01312 | - | - | - |
| | b | - | - | - | - | - | - | DDT19_01310 | - | - | - |
| | c | - | - | - | - | - | - | DDT19_01311 | - | - | - |
| | c | - | - | - | - | - | - | - | - | - | - |

**Table S4. Environmental paramers and chemical coposition of Hakuba Happo spring water**

| | July 2016 | October 2016 | October 2017 |
|---|---|---|---|
| pH | 10.95 | 10.80 | 10.67 |
| Temperature (ºC) | 47.5 | 47.4 | 45.6 |
| ORP (mV) | -435 | -432 | -453 |
| electrical conductivity (mS/m) | 47.7 | 43.6 | 51.7 |
| dissolved oxygen (mg/L) | <dl | <dl | <dl |
| $Na^+$ (ppm) | 32 | 30 | <dl |
| $K^+$ (ppm) | <dl | <dl | <dl |
| $Ca^{2+}$ (ppm) | <dl | <dl | <dl |
| $NO_3^-$ (ppm) | <dl | <dl | <dl |
| $NH_4^+$ (µM) | ND | ND | 2.9 |
| Amino acids | | | |
| Aspartic Acid (nM) | ND | ND | <dl |
| Threonine (nM) | ND | ND | <dl |
| Serine (nM) | ND | ND | <dl |
| Glutamic Acid (nM) | ND | ND | <dl |
| Glycine (nM) | ND | ND | 5.4 ± 1.6 |
| Alanine (nM) | ND | ND | <dl |
| Cysteine (nM) | ND | ND | <dl |
| Valine (nM) | ND | ND | <dl |
| Methionine (nM) | ND | ND | <dl |
| Isoleucine (nM) | ND | ND | <dl |
| Leucine (nM) | ND | ND | <dl |
| Tyrosine (nM) | ND | ND | <dl |
| Phenylalanine (nM) | ND | ND | <dl |
| Histidine (nM) | ND | ND | <dl |
| Lysine (nM) | ND | ND | <dl |
| Arginine (nM) | ND | ND | <dl |

<dl, below our quantification limit; ND, not determined
